# Engineering of a biosensor for intracellular aspartate

**DOI:** 10.1101/2023.05.04.537313

**Authors:** Lars Hellweg, Martin Pfeifer, Lena Chang, Miroslaw Tarnawski, Andrea Bergner, Jana Kress, Julien Hiblot, Jürgen Reinhardt, Kai Johnsson, Philipp Leippe

## Abstract

Aspartate is a limiting metabolite in proliferating cells with its production closely linked to glutamine and mitochondrial metabolism. To date, measuring aspartate concentrations in live cells was deemed impossible. We present iAspSnFR, a genetically-encoded biosensor for intracellular aspartate, engineered by displaying and screening biosensor libraries in HEK293 cells. In live cells, iAspSnFR exhibits a dynamic range of 130% fluorescence change and detects reduced aspartate levels upon glutamine deprivation or glutaminase inhibition. Furthermore, iAspSnFR tracks aspartate uptake by excitatory amino acid transporters, or of asparagine after co-expression of an asparaginase. Importantly, iAspSnFR reports aspartate depletions upon electron transport chain inhibition, and therefore it can serve as a proxy for mitochondrial respiration. Consequently, iAspSnFR can dissect the major cellular pathways of aspartate production, offering immediate applications, particularly in cancer biology, such as screening small molecules targeting aspartate and glutamine metabolism.

## Introduction

Aspartate plays a vital role in cellular metabolism, including protein and nucleotide synthesis, and ammonia detoxification^1^. For example, in proliferating cells, aspartate is essential for the synthesis of purine and pyrimidine nucleotides, which are required for DNA replication and RNA transcription^2,3^. In respiring cells, aspartate synthesis occurs in mitochondria by transamination of the tricarboxylic acid cycle (TCA cycle) intermediate oxaloacetate (OAA), closely linking aspartate to mitochondrial metabolism. Glutamine can serve as an anaplerotic source for aspartate production after its conversion to glutamate by glutaminase (**Fig. 1a**)^4^. Aspartate is exported to the cytoplasm by aspartate-glutamate-carriers (AGC1/2, or SLC25A12/13), which exchange aspartate for glutamate plus a proton^5^. Along with the malate-ketoglutarate carrier SLC25A11, AGCs form the malate-aspartate shuttle that transfers cytoplasmic reducing equivalents, such as those derived from glycolysis, to the respiratory chain (RC)^6^. Maintaining aspartate levels is critical for cell proliferation and survival, particularly in cells with compromised electron transport chain function, a common trait in cancer cells^2,3^. This metabolic vulnerability is also present *in vivo*, emphasizing the importance of aspartate in cellular metabolism and growth^7^.

**Fig. 1.**
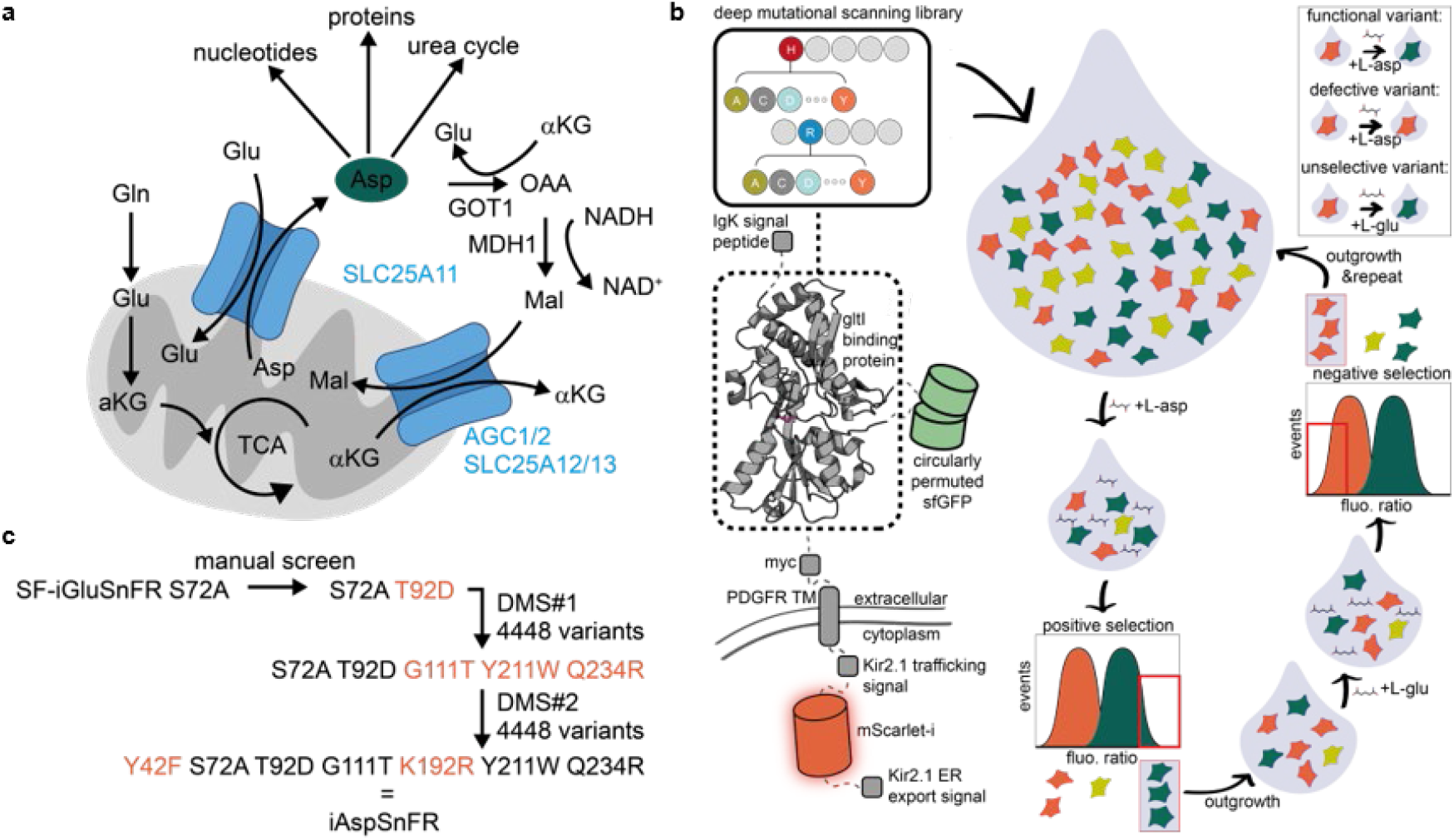
Engineering iAspSnFR through cell surface display and screening. **a)** Overview of the multiple roles of aspartate in cellular and mitochondrial (grey) metabolism. **b)** Scheme of the screening strategy to engineer iAspSnFR. SF-iGluSnFR-S72A (green: SF-GFP, grey: glutamate/aspartate periplasmic binding protein) was modified to express mScarlet-I (red) intracellularly for normalization. Addition of aspartate to cells increases the reporter sfGFP emission and thus sfGFP/mScarlet-I emission ratio (fluo. ratio). A deep mutational scanning (DMS) library of the gltI aspartate/glutamate binding protein domain was generated and delivered to HEK293-JI cells with the R4 integrase system so that only one variant is expressed per cell. Cells were placed in suspension and exposed to aspartate, after which only those expressing functional variants exhibited an increase in fluo. ratio. Those cells were then enriched using fluorescence-activated cell sorting (FACS) and subsequently outgrown. Sanger or next-generation sequencing (NGS) was used to analyze enrichments before proceeding to a second round of enrichment. In this step, glutamate was added, and only those cells that maintained a low fluo. ratio indicating selectivity for aspartate were isolated. This process is repeated over multiple rounds. **c)** iAspSnFRs protein engineering lineage.

Measurements of aspartate concentrations in cells have primarily relied on mass spectrometry (MS). Yet MS-based methods, while powerful, can be limited in throughput due to sample preparation, require complex experimental workflows for organellar fractionation or absolute quantification^8^, and only provide terminal end-point measurements. Genetically-encoded fluorescent biosensors offer an ideal complementary solution to MS, as they are compatible with time-resolved measurements in live cells, adaptable to high throughput settings and can be genetically targeted to organelles.

In this study, we present iAspSnFR, a genetically-encoded, green fluorescent biosensor for intracellular aspartate. Engineered through deep mutational scanning libraries displayed and screened on mammalian cells, iAspSnFR is specific for aspartate and bright with 1000% fluorescence increase *in vitro*. In live cells, we demonstrate that iAspSnFR is capable of detecting both increases and decreases of cytoplasmic aspartate over a range of perturbations targeting aspartate uptake or metabolism.

## Results

### Engineering of aspartate sensor iAspSnFR

We began by exploring the rich history of glutamate sensors based on the periplasmic aspartate/glutamate binding protein gltI^9^, which dates back about two decades^10–17^. Wild-type gltI exhibits a low micromolar affinity both for aspartate (K_D_ = 0.7 μM) and glutamate (K_D_ = 1.2 μM)^9^ and has been used for the construction of fluorescence resonance energy transfer (FRET) sensors^11–13^ and intensiometric sensors based on circularly permuted GFP^14,15^ or superfolder-GFP^16,17^. We titrated some of the intensiometric sensors as purified proteins against aspartate and glutamate. Purified SF-iGluSnFR-S72A^16^ was found to bind aspartate with K_0.5_ = 11 μM, glutamate with K_0.5_ = 0.38 mM and asparagine with K_0.5_ = 4.4 mM (**Supplementary Fig. 1**). Therefore, SF-iGluSnFR-S72A already exhibits a desirable selectivity for aspartate, albeit with much too high affinity. Based on this scaffold, we set out to identify mutations that would lower affinity to the intracellular aspartate range (∼0.1 – 10 mM^8^) while remaining insensitive to glutamate, the most abundant cytoplasmic amino acid present at approximately 20 mM^8^. For normalization, SF-iGluSnFR-S72A was modified to express the red fluorescent protein mScarlet-I at the intracellular *C*-terminus (**Fig. 1b**), yielding fluorescence ratio (hereafter abbreviated as fluo. ratio) by dividing sfGFP and mScarlet-I fluorescence intensity units. Then, saturation mutagenesis was individually performed on residues E25, S72, T92 and V184, all positions previously reported to affect selectivity and/or affinity^13,15,16^. Those variants were cloned individually, and displayed one-by-one on the surface of HeLa cells by transient transfection. Increasing concentrations of aspartate (0.1, 1, 10 mM) were perfused while monitoring fluorescence by widefield microscopy to measure affinity, followed by a high concentration of glutamate (10 mM) to assess selectivity (**Supplementary Fig. 2**). From this screen, SF-iGluSnFR-S72A T92D was identified as the best candidate, based on its ∼1 mM aspartate affinity, acceptable ∼400% dynamic range, and no detectable binding to glutamate up to 100 mM (**Supplementary Fig. 3a,b**).

However, when stably expressing SF-iGluSnFR-S72A T92D as soluble protein in the cytoplasm of HEK293 Jump In™ T-Rex™ cells (hereafter abbreviated as HEK293-JI) and titrating against aspartate, an increase in fluorescence could not be observed (**Supplementary Fig. 3c**). We went back and titrated the starting template SF-iGluSnFR-S72A in a similar fashion. In HEK293-JI cell lysate, SF-iGluSnFR-S72A responded with approximately 70% fluorescence increase to aspartate with K_0.5_ = 16 μM (10-24 μM 95%CI) and to glutamate with K_0.5_ = 0.50 mM (0.43-0.59 mM 95% CI) (**Supplementary Fig. 3d**). As such, while the dynamic range was decreased in lysate, the affinity values of SF-iGluSnFR-S72A were very similar to purified protein (**Supplementary Fig. 1d**), ruling out that SF-iGluSnFR variant expression was generally incompatible with the cytoplasm. Due to the favorable selectivity profile of SF-iGluSnFR-S72A T92D on the cell surface, we decided to use it as new scaffold and search for allosteric mutations stabilizing its non-fluorescent and fluorescent conformational states, as assessed by absolute fluorescence in presence of aspartate and dynamic range. To do this holistically and increase the likelihood of identifying such beneficial mutations at novel positions, we performed a deep mutational scan (DMS)^18^ (**Fig. 1b, box**) over the entire large fragment of the gltI binding protein (residues 8-255 = 247 amino acids), yielding a biosensor library of 4448 variants (**Supplementary Data 1**). To increase throughput, we decided to screen in a pooled fashion, where the biosensor variant library is introduced to a single population of cells. This population is then enriched for improved variants, and sequenced to reveal the underlying sequence identity. The enrichment strategy was based on the observation that HEK293-JI cells expressing cell surface SF-iGluSnFR-S72A T92D produced robust fluorescent increases when taken in suspension, mixed with saturating aspartate and analyzed with flow cytometry. This indicated that only a limited dilution of sample fluid and shear fluid to form droplets occurs within the flow cytometer. By comparing titrations of adherent live cells at the plate reader and suspended cells in the flow cytometer, this dilution factor was experimentally determined to be 5-fold (**Supplementary Fig. 4**). Therefore, we reasoned that by displaying a library of biosensors on the cell surface and exposing the library to saturating aspartate, fluorescence-activated cell sorting (FACS) could be used to enrich for variants with better properties, *i*.*e*. higher absolute fluorescence (positive selection, **Fig. 1b**). Additionally, this provided a robust way to screen for selectivity by exposure of cells to glutamate and enriching cells that remained in a low fluorescent state (negative selection). For library delivery, the commercially available HEK293-JI cell system using the R4 integrase was chosen. When delivering a library of donor plasmids to a population of HEK293-JI cells, only a single variant per cell will be integrated since only one R4 *attP* site exists per cell genome. After DMS library integration into HEK293-JI cells, enrichments were performed in four subsequent rounds of enrichment and outgrowth, collecting the best performing 5-15% of cells from each round. Populations of each round were analyzed by next-generation sequencing (**Supplementary Fig. 5**). After the fourth round, two variants were enriched to a very high abundancy, with G111T present at 32% variant frequency (counts of variant reads / all reads) and Q234R at 31%, followed by N93R at 9% and Y211W at 5%. The two highly frequent G111T and Q234R mutations were selected for validation, with mutation Y211W, located at the back of the binding pocket and hypothesized to tighten binding of the aspartate ammonium group. To verify screening hits, the dynamic range and absolute brightness of variants G111T, Q234R and Y211W individually integrated in HEK293-JI cells were measured, confirming performance improvements for all three mutations (**Supplementary Fig. 6a**). Next, cleared lysate of HEK293-JI cells expressing the cytoplasmic DMS variants were titrated against aspartate and glutamate. All mutations individually recovered sensor performance, restoring K_0.5_ to 9-23 mM and dynamic range to 50-160% (**Supplementary Fig. 6b-e**). Interestingly, the combination of all DMS mutations was synergistic for increasing the dynamic range, yielding the triple mutant G111T Y211W Q234R with 25 mM affinity and 310% dynamic range (**Supplementary Fig. 6f)**. This prompted us to repeat the DMS with increased stringency (top 1% cells) over only two rounds of positive selection, resulting in the additional K192R and Y42F mutations. Those two additional mutations were incorporated into the design, finally yielding a mutant with a total of 6 mutations (Y42F T92D G111T K192R Y211W Q23R) compared to SF-iGluSnFR-S72A (**Fig. 1c**). When expressed in the cytoplasm, and titrated against aspartate in cleared lysate, this mutant responded with a 640% change in fluo. ratio at 1.4 mM apparent affinity, while remaining insensitive to glutamate up to 95 mM (**Supplementary Fig. 6g, h)**, meeting our initial engineering goals. Accordingly, this mutant was termed iAspSnFR and was subjected to in-depth *in vitro* characterization.

### *In vitro* characterization of iAspSnFR

To understand the respective specificities of aspartate and glutamate sensors, the X-ray structures of apo- and glutamate-bound SF-iGluSnFR-S72A, as well as aspartate-bound iAspSnFR, were obtained at 2.6 Å, 1.7 Å and 1.7 Å (**Fig. 2a, Table S3**). Noteworthy, a citrate molecule was observed in the gltI aspartate/glutamate binding pocket of apo*-*SF-iGluSnFR-S72A, which was present at very high concentration (1.5 M) in the crystallization condition (**Supplementary Fig. 7**). Competition titrations with citrate were performed and ruled out any specific binding and interference with sensor performance (**Supplementary Fig. 8**). iAspSnFR’s aspartate-sensitive fluorescence (Ex. 490 nm, Em. 512 nm) is normalized to the mScarlet-I fluorescence (Ex. 570 nm, Em. 594 nm; **Fig. 2b, d**). Purified iAspSnFR binds aspartate with a high 1000% fluorescence increase and an apparent affinity of 0.9 mM (95% CI: 0.74-1.2 mM). It demonstrates 52-fold selectivity for aspartate over glutamate and 35-fold selectivity for aspartate over asparagine (**Fig. 2c**). iAspSnFR is insensitive to pH fluctuations between pH = 7-9, but sensitive to acidic pH with pKa ≈ 6 in the aspartate bound state (**Fig. 2e**). iAspSnFR was titrated with aspartate in the presence of competing amino acids or drugs to evaluate competitive effects (**Fig. 2f**). At 5 mM, asparagine was found to interfere slightly with the titration. To further investigate this effect, aspartate was titrated in presence of varying concentrations of competing asparagine (**Supplementary Fig. 9a**). Similarly, competing glutamate concentrations were increased stepwise to 20 mM (**Supplementary Fig. 9b**). Based on those titrations, iAspSnFR measurements are not affected by glutamate up to 20 mM but asparagine can interfere with the measurement when aspartate is low (<100 μM) and asparagine high (>3 mM). However, at physiological concentrations, asparagine can only elicit a small fraction of the fluorescence response compared to aspartate (9.5 mM asparagine: 1.6 norm. fluo. ratio; 9.5mM aspartate: 9.7 norm. fluo. ratio, **Fig. 2c**). Furthermore, a whole-cell ratio of 3.6 aspartate to 1 asparagine was measured by mass spectrometry in control HEK293-JI GFP-OE cells (**Supplementary Fig. 10**). In combination, while caution should be exercised in special cases where aspartate is expected to be low (<100 μM) and asparagine high (>3 mM), asparagine interference should be of minimal concern for experiments in cells. Lastly, iAspSnFR’s dynamic range was found to be negatively impacted by temperatures above 30°C (**Supplementary Fig. 11**), likely introduced during performing the DMS enrichments with FACS at room temperature. Subsequent cell experiments were thus measured at 30°C on the microscope or at room temperature for flow cytometry analysis to ensure maximum iAspSnFR dynamic range.

**Fig. 2.**
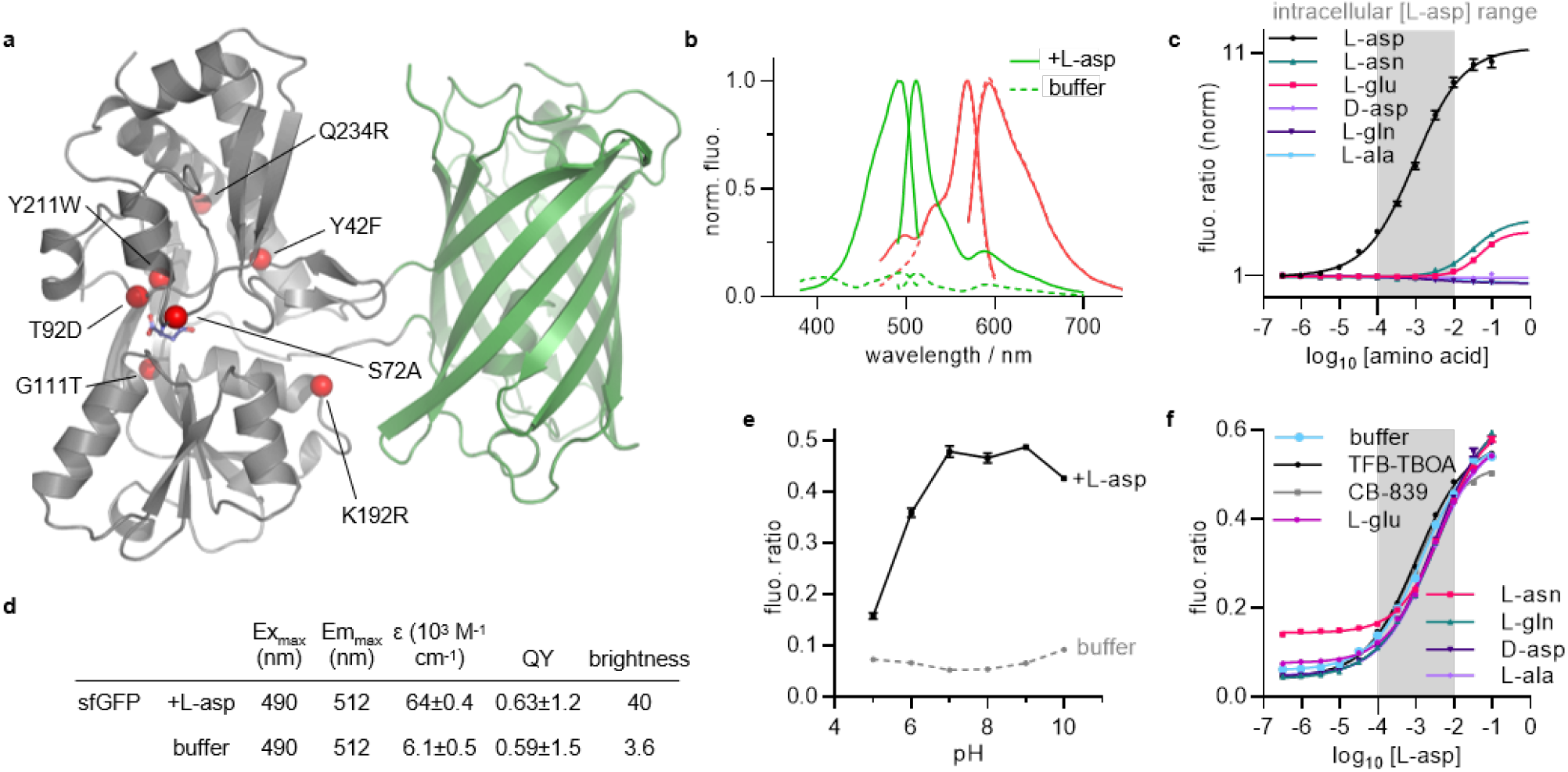
*In vitro* characterization of iAspSnFR. **a)** Crystal structure of iAspSnFR bound to L-asp. Red spheres indicate mutations compared to SF-iGluSnFR (grey: gltI periplasmic aspartate/glutamate binding protein, green: sfGFP, blue sticks: aspartate). **b)** Excitation and emission spectra in presence or absence of L-asp (green: sfGFP, red: mScarlet-I). **c)** Titration curve for aspartate and related amino acids. K_0.5_ (95% CI): aspartate = 0.9 mM (0.74-1.2 mM), asparagine = 32 mM (0.02-0.27 M), glutamate = 47 mM (36-76 mM). Mean±SEM **d)** Spectroscopic properties (Ex: Excitation, Em: Emission, ε: Extinction coefficient, QY: Quantum yield). Mean±SD. **e)** pH-sensitivity. Mean±SD **f)** Titration curve with aspartate in presence of competing amino acids and chemical probes. (Concentrations: amino acids 5 mM, TFB-TBOA 10 μM, CB-839 10 μM). Mean±SD. All titrations were performed in technical quadruplicates.

### iAspSnFR detects perturbations of cytoplasmic aspartate in HEK293-JI cells

To test if iAspSnFR can detect perturbations of aspartate metabolism in live cells, cells were first subjected to nutrient starvation for 3 hrs. This decreased intracellular aspartate, as evidenced by a reduced fluo. ratio to about 0.7-fold of basal, which could be restored to above basal (1.1-fold) upon addition of high external 10 mM aspartate (**Fig. 3a, b, j**). To further increase intracellular aspartate and fully saturate iAspSnFR, HEK293-JI cells overexpressing the glutamate/aspartate transporter SLC1A3 and iAspSnFR (**Fig. 3c**) were examined. Cells were monitored over 5 minutes before addition of 1 mM aspartate, which led to a gradual increase in fluo. ratio that plateaued to 1.5-fold after 15 min, consistent with direct aspartate uptake by SLC1A3 (**Fig. 3d**). For quantification, the experiment was repeated, measuring steady-state aspartate levels (**Fig. 3e**). Incubation with 0.5 mM aspartate increased the fluo. ratio to 1.7-fold compared to starvation, or to 2.1-fold when compared to cells treated with aspartate and the SLC1A3 inhibitor TFB-TBOA (**Fig. 3e**). Notably, starving SLC1A3-overexpressing cells did not reduce intracellular aspartate (fluo. ratio = 0.27) to levels similar of non-expressing cells (**Fig. 3b**, fluo. ratio = 0.22). This suggests that SLC1A3 takes up aspartate from the medium, potentially from dying or aspartate secreting cells^19^. To determine the K_0.5_ of SLC1A3 for aspartate, the experiment was repeated using a full aspartate dose-response concentration series, and analyzed by flow cytometry (**Fig. 3f**). This revealed that aspartate uptake proceeds in two phases, with the first phase following a sigmoid dose-response curve with K_0.5_ = 9-47 μM (95% CI) that was inhibited by TFB-TBOA, consistent with uptake mediated by SLC1A3. The second phase at high, supraphysiological external aspartate concentrations (>10 mM) was not inhibited by TFB-TBOA, suggesting passive diffusion or aspartate “hitchhiking” through other endogenously expressed transporters^20^.

Next, we focused on SLC1A5/ASCT2, which catalyzes the antiport of a range of neutral amino acids, such as glutamine, alanine, serine, threonine and asparagine^21^. SLC1A5 is highly expressed in a variety of cancers^22^, making it an attractive drug discovery target. However, SLC1A5 assays rely on electrophysiological recordings or radioactive tracers, not amenable to high throughput screens. We therefore asked if we could use iAspSnFR to develop a fluorescence-based assay and readout for amino acid uptake by SLC1A5. Indeed, asparagine uptake in HEK293-JI cells should primarily be mediated by SLC1A5, since proteomics and metabolomics analyses of asparagine uptake transporters indicated SLC1A5 to be by far the highest expressed asparagine transporter in HEK293-JI cells (**Supplementary Fig. 12**). To readout asparagine uptake with iAspSnFR, the *guinea pig* asparaginase (gpASNase1)^7,23^, which converts asparagine to aspartate, was co-expressed in HEK293-iAspSnFR cells. Indeed, incubation with 0.5 mM asparagine increased aspartate levels in gpASNase1-transfected cells only (**Fig. 3g, j**). This increase could be blocked by addition of a 10 mM alanine, which competes with asparagine for uptake. The same experiment was repeated with a full concentration series and analyzed by flow cytometry, revealing a K_0.5_ = 20-34 μM (95% CI) for asparagine uptake and its conversion to aspartate (**Fig. 3h**). Finally, we examined endogenous aspartate production from oxidative glutamine metabolism (**Fig. 3i, k**). HEK293-iAspSnFR cells were deprived of nutrients to deplete aspartate, which could be restored by addition of glutamine alone. In glutamine containing medium, incubation with the glutaminase inhibitor CB-839 alone was sufficient to mimic nutrient deprivation. To test if cytoplasmic aspartate levels can readout respiratory chain (RC) activity, the RC inhibitors rotenone or antimycin A were added in presence of glutamine. RC inhibition fully halted aspartate production, indicating that aspartate production from glutamine occurs in mitochondria in dependence of RC activity. Pyruvate addition could partially rescue aspartate levels, presumably by serving as substrate for pyruvate carboxylase, and as exogenous electron acceptor^3^ to partially restore the NAD^+^/NADH imbalance caused by RC inhibition. As such, the readout of cytoplasmic aspartate levels by iAspSnFR can serve as a fluorescent proxy of redox state and mitochondrial respiration.

**Fig. 3.**
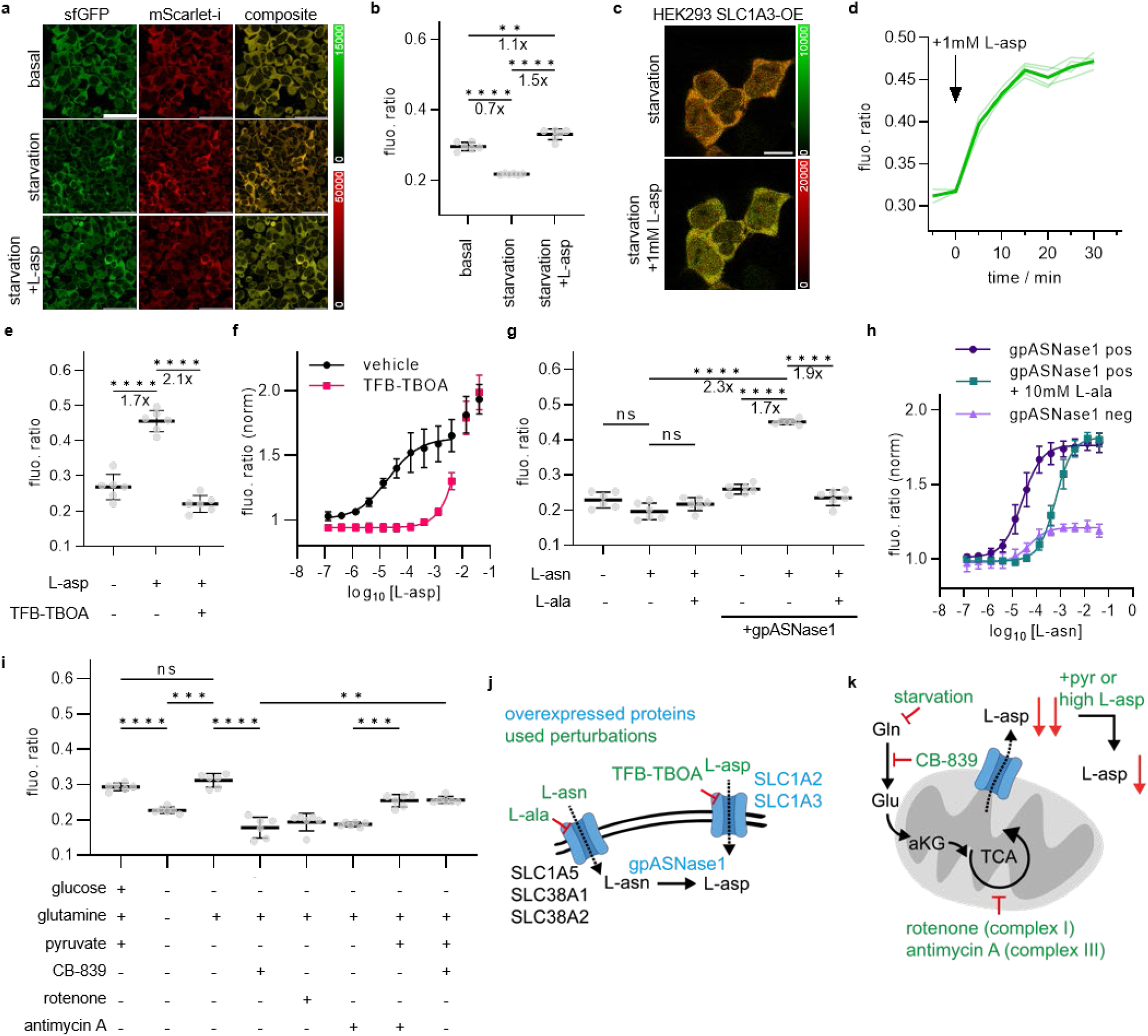
iAspSnFR detects perturbations of cytoplasmic L-asp in HEK293-JI cells. **a)** Confocal images of HEK293-JI-iAspSnR cells after 3 hrs nutrient starvation, including a composite of the sfGFP channel (green) and the mScarlet-I channel (red). The fluo. ratio of HEK293-JI-iAspSnR cells is reduced upon nutrient starvation and restored to above basal after addition of 10 mM aspartate (scale bar = 50 μm, steady state after 3 hrs incubation). **b)** Quantitative analysis of experiments presented in a). Fluo. ratio is obtained by dividing sfGFP by mScarlet-I fluorescence intensity values. (Mean±SD of six wells per condition with 65-169 cells each over two independent experiments). **c)** Composite of confocal images (scale bar = 10 μm, green: sfGFP, red: mScarlet-I) of HEK293-JI-SLC1A3 cells transiently transfected with iAspSnFR. Top: before aspartate addition, bottom: after aspartate addition at t=30 min. **d)** Full timecourse of the 4 cells in c). **e)** Quantitative analysis of experiments presented in c) and d) after treatment for 3 hrs. (L-asp = 0.5 mM + 0.04% DMSO, TFB-TBOA = 10 μM; all in starvation medium; mean±SD, six wells with 10-45 cells each over two independent experiments). **f)** Same experiment as c) but measured with flow cytometry. HEK293-JI-SCL1A3 cells transiently transfected with iAspSnFR were starved for 3 hrs, taken into suspension and incubated with the indicated compounds for 30 min before flow cytometry analysis. (K_0.5_(L-asp) = 9-47 μM (95% CI); mean±SD of n = 4 replicates over 2 independent experiments, each replicate represents ∼1400-3000 cells; vehicle = 0.04% DMSO, TFBO-TBOA = 10 μM). A non-linear component of aspartate uptake was observed at high concentrations (>10 mM L-asp) that couldn’t be blocked by TFB-TBOA. **g)** HEK293-JI-iAspSnR were transiently transfected with EBFP-gpASNase1. Incubation with 0.5 mM asparagine increases cytoplasmic aspartate only in EBFP-gpASNase1 transfected cells. Competition for asparagine uptake by 10 mM alanine blocked this effect. All conditions in starvation medium. **h)** HEK293-JI-iAspSnR were transiently transfected with EBFP-gpASNase1 and nutrient starved for 3 hrs, then incubated with varying concentrations of asparagine in absence or presence of 10 mM alanine. The sample was gated for EBFP-gpASNase1 expression and analyzed for iAspSnFR fluo. ratio. K_0.5_(L-asn) = 20-34 μM (95% CI), K_0.5_(L-asn + 10 mM L-ala) = 0.54-0.74 mM (95% CI); mean±SD of n = 4 replicates over 2 independent experiments, each replicate represents ∼1000-11000 cells. **i)** Pharmacological perturbations of steady-state aspartate levels in HEK293-JI-iAspSnFR cells. Glutamine withdrawal, glutaminase inhibition by CB-839 or inhibition of the electron transport chain (antimycin A, rotenone) deplete cytoplasmic aspartate levels. Pyruvate addition partially restores aspartate levels. (Mean±SD of 6 wells per condition with 13-204 cells each over two independent experiments). **j)** Overview of experiments targeting asparagine or aspartate uptake in c-h). **k)** Overview of steady state perturbation of aspartate metabolism in a,b,i). Significance testing: Welch ANOVA tests with Dunnett’s T3 multiple comparison correction. *P*-values: 0.1234 (ns), 0.0332 (*), 0.0021 (**), 0.0002 (***), <0.0001 (****).

## Discussion

In this study, we introduced iAspSnFR, the first genetically-encoded biosensor for intracellular aspartate. Our innovative cell surface display and screening technology to engineer iAspSnFR combines deep mutational scanning, mammalian cell surface display, fluorescence-activated cell sorting, and next-generation sequencing. We plan to further refine our technology to screen biosensor libraries directly in the cytoplasm or mitochondria. Future generations of iAspSnFR will aim to address temperature sensitivity, increase dynamic range in cells, and target organelles, particularly mitochondria.

We thoroughly characterized iAspSnFR, demonstrating its ability to accurately measure aspartate concentrations with minimal effects from competing amino acids or drugs. Cell experiments showed that iAspSnFR can reliably detect both increases and decreases in cytoplasmic aspartate in response to perturbations of amino acid uptake, metabolism, and mitochondrial function.

Aspartate plays a critical role in amino acid synthesis, nucleotide synthesis^18^, the urea cycle^24^ and redox homeostasis^6^. Consequently, aspartate metabolism is implicated in in various pathological contexts, such as cancer cell proliferation^2,3,25,26^, stem cell differentiation^27,28^ and urea cycle disorders^29,30^. By providing a means to monitor intracellular aspartate levels in live cells, iAspSnFR sets the stage for mechanistic investigations into the underlying defects in aspartate metabolism and may help identify therapeutic interventions.

## Methods

See Supplementary Information.

## Data and materials availability

The iAspSnFR plasmid is deposited and distributed at addgene with accession number 201394 (pcDNA3.1/Neo). The X-ray crystal structures of SF-iGluSnFR-S72A, SF-iGluSnFR-S72A in complex with L-aspartate and iAspSnFR in complex with L-aspartate were deposited to the PDB with accession codes 8OVN, 8OVO and 8OVP, respectively. Source data are provided with the article. Included RESOLUTE datasets are publicly available at www.re-solute.eu/datasets. Protocols for their acquisition are publicly available at https://re-solute.eu/resources/reagents. Additional materials or data are available by inquiry to the corresponding authors upon reasonable request.

## Acknowledgements

We thank Julian Kompa for measuring quantum yields, Dr. Birgit Koch for assistance in cell culture, and Dr. Sebastian Fabritz with the mass spectrometry core facility for MS measurements. Diffraction data were collected at the Swiss Light Source, beamline X10SA, of the Paul Scherrer Institute, Villigen, Switzerland. PL is grateful to Dr. Klaus Seuwen for inspiration and support. We acknowledge the RESOLUTE consortium for providing reagents and datasets, and thank all consortium colleagues for critical feedback.

This study received funding from the Max Planck Society, from the Deutsche Forschungsgemeinschaft (DFG, German Research Foundation) TRR 186, and is part of the RESOLUTE consortium. RESOLUTE has received funding from the Innovative Medicines Initiative 2 Joint Undertaking under grant agreement No 777372. This Joint Undertaking receives support from the European Union’s Horizon 2020 research and innovation programme and EFPIA. This article reflects only the authors’ views and neither IMI nor the European Union and EFPIA are responsible for any use that may be made of the information contained therein.

---

**Supplementary Fig. 1.**
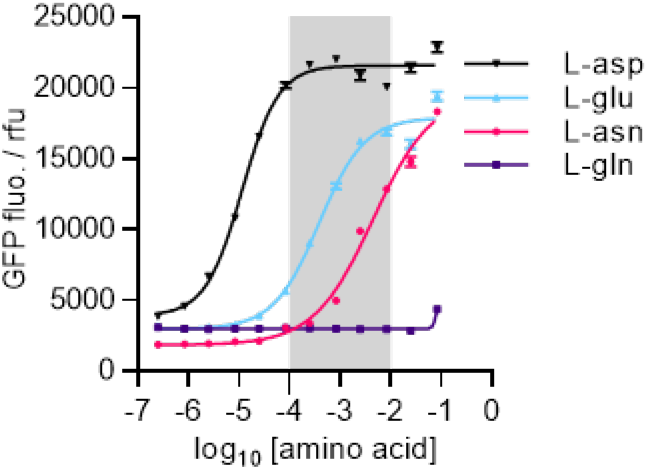
Titration of SF-iGluSnFR-S72A expressed in *E. Coli*. and purified. Mean±SD of quadruplicates. K_0.5_ values (95% CI): L-asp = 11 μM (9.8-13 μM), L-glu = 0.38 mM (0.31-0.47 mM), L-asn = 4.4 mM (3.4-6.4 mM), L-gln = N/A.

**Supplementary Fig. 2.**
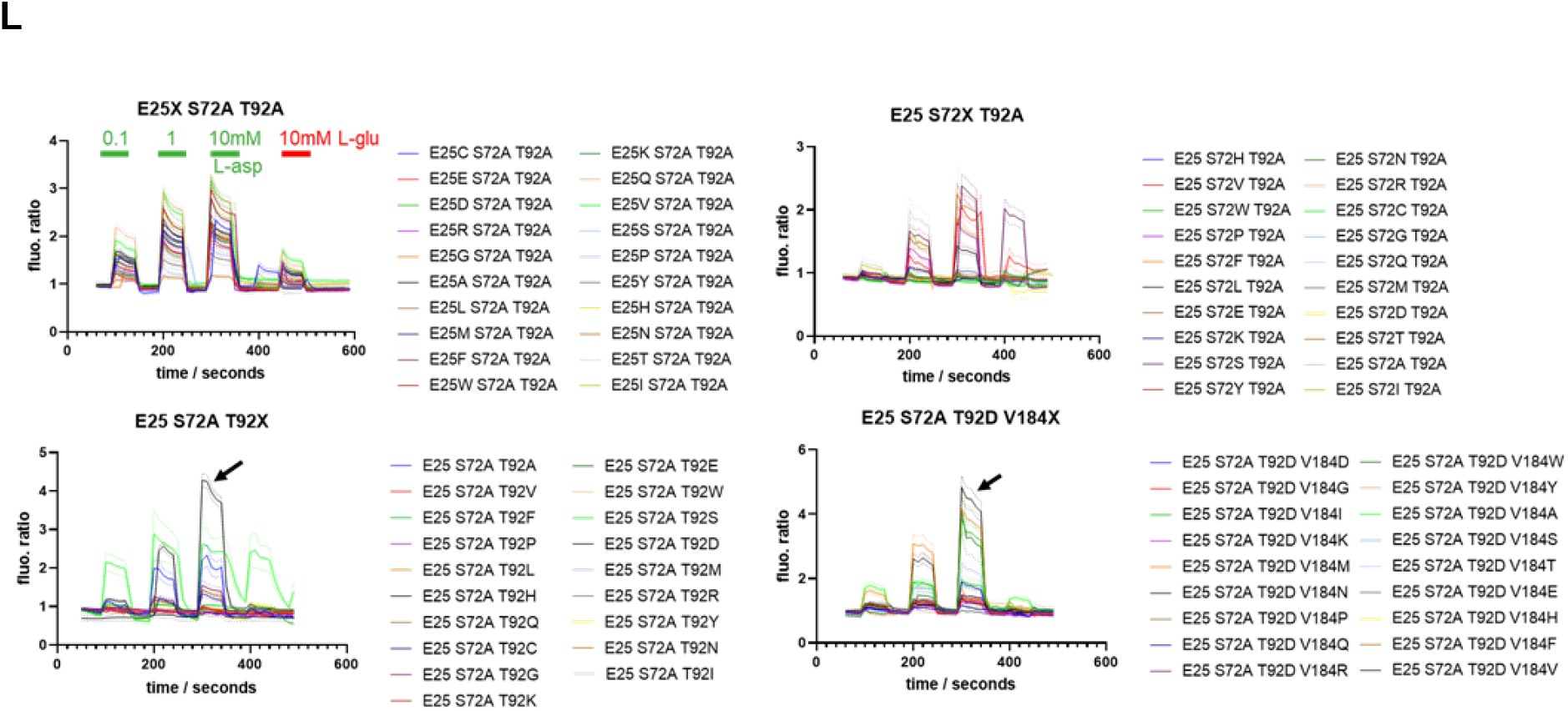
Manual one-by-one perfusion experiments in HeLa cells transiently with SF-iGluSnFR variants. The most promising candidate SF-iGluSnFR-S72A T92D is marked with a black arrow.

**Supplementary Fig. 3.**
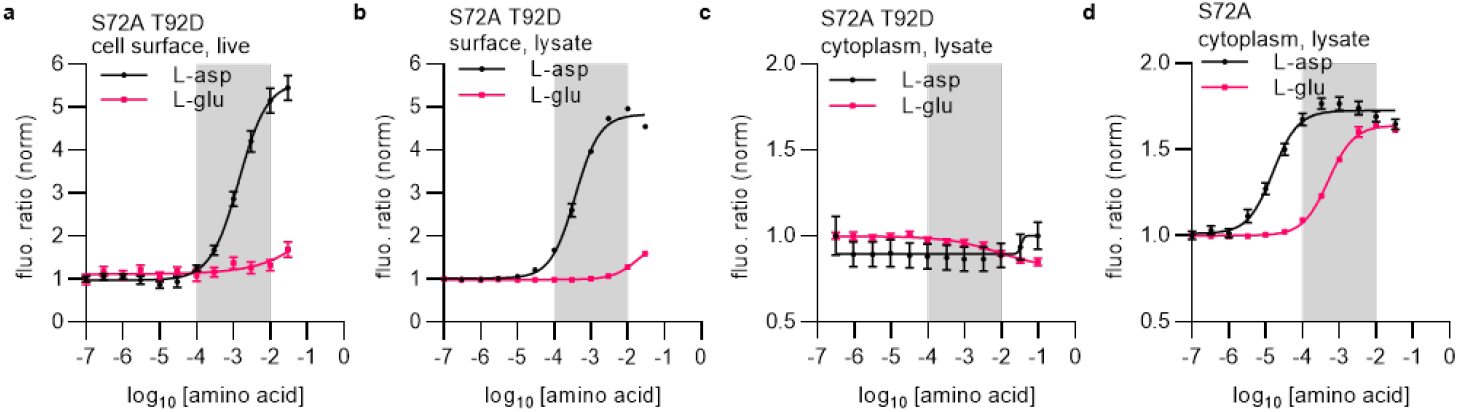
**a)** Cell surface SF-iGluSnFR-S72A T92D in live HEK293-JI cells, measured at the plate reader. **b)** Same as a) but using cleared cell lysate (to exclude effects from the lysis protocol) **c)** titration in cleared lysate of cyto-SF-iGluSnFR-S72A T92D or **d)** cyto-SF-iGluSnFR-S72A expressing HEK293-JI cells. Titrations in cleared lysate should be considered a lower boundary for sensor performance since the protein/lysate solution is crude, and small molecules or autofluorescent (NAD(P)H, FAD, …) cell components from lysate preparation (i.e. also aspartate and glutamate from cells) are still present. Grey box: intracellular aspartate range.

**Supplementary Fig. 4.**
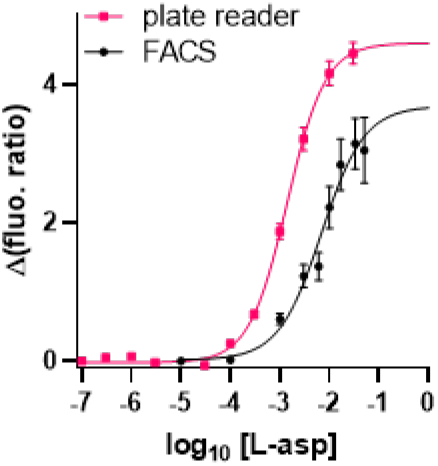
Quantification of the dilution factor within the FACS instrument by comparing the titrations of HEK293-JI expressing cell-surface SF-iGluSnFR-S72A T92D as adherent cells at the platereader (pink) and at the FACS instrument (black). The black dots are the nominal concentrations that are added to the sample cell suspension. Cells are exposed to a lower than nominal concentration at the point of measurement within the FACS instrument due to dilution of sample fluid with shear fluid. The K_0.5_ measured at the FACS is approx. 5-times lower than on live cells, indicating an ∼5-fold dilution factor.

**Supplementary Fig. 5.**
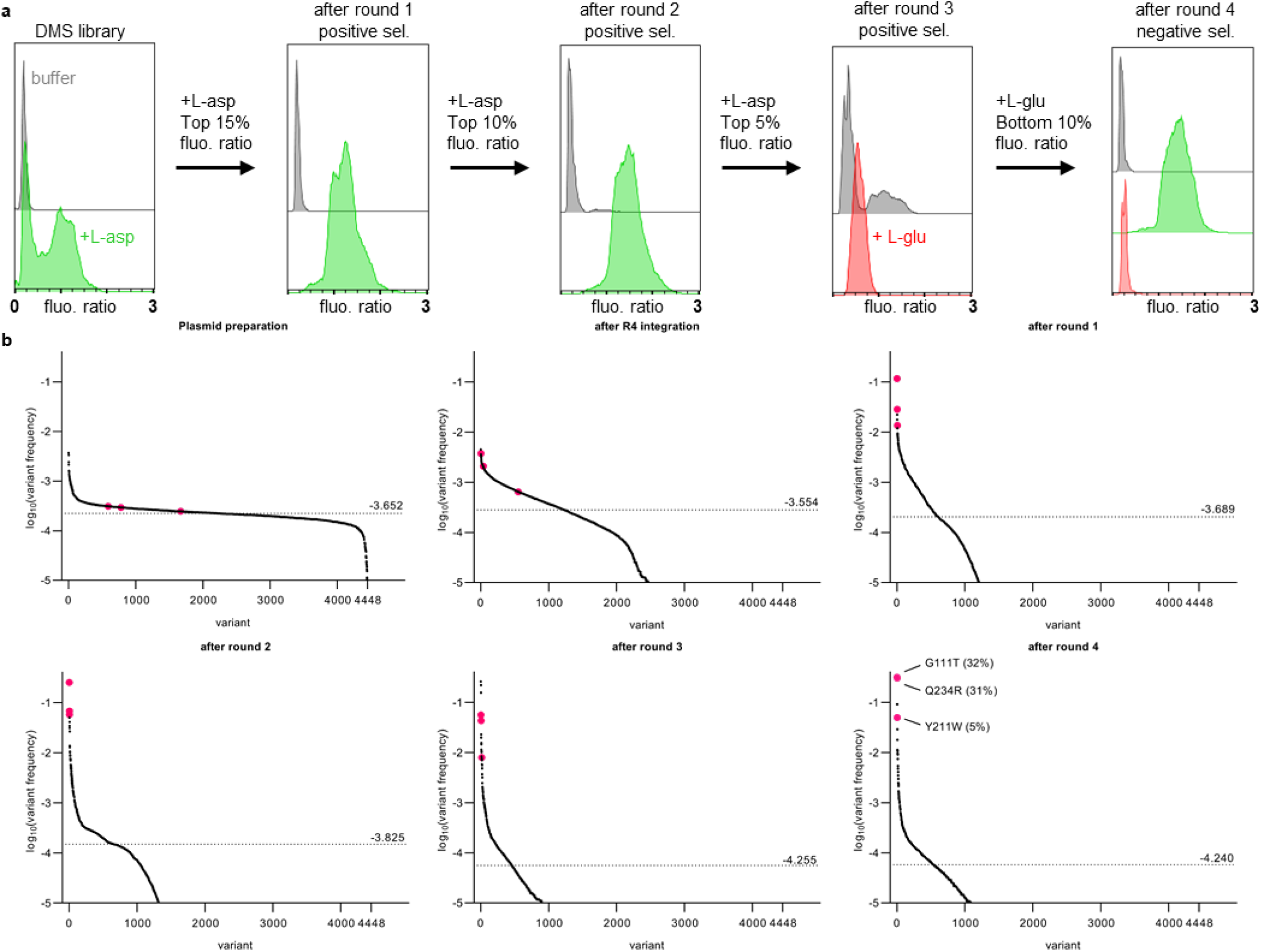
DMS enrichments and NGS Analysis. **a)** Fluo. ratio of the initial cell populations of DMS#1 and after each selection round. **b)** Variant frequency (variant-reads / all reads) as analysed by Illumina sequencing, dotted line at median. Comparison of the plasmid library with the library after R4 integration revealed a bottleneck, resulting in a skewed library. The library diversity is decreasing rapidly after selection round 1, as about half of the variants produce non-functional sensor (see leftmost panel of a), dim cells after aspartate addition). The three final screening hits (G111T Y211W and Q234R) which were analysed in depth after round 4 (pink dots) emerge rapidly. In the final cell population, these three screening hits together comprise 68% of all observed sequences. Dotted line at median frequency (variant cutoff at 10^−5^).

**Supplementary Fig. 6.**
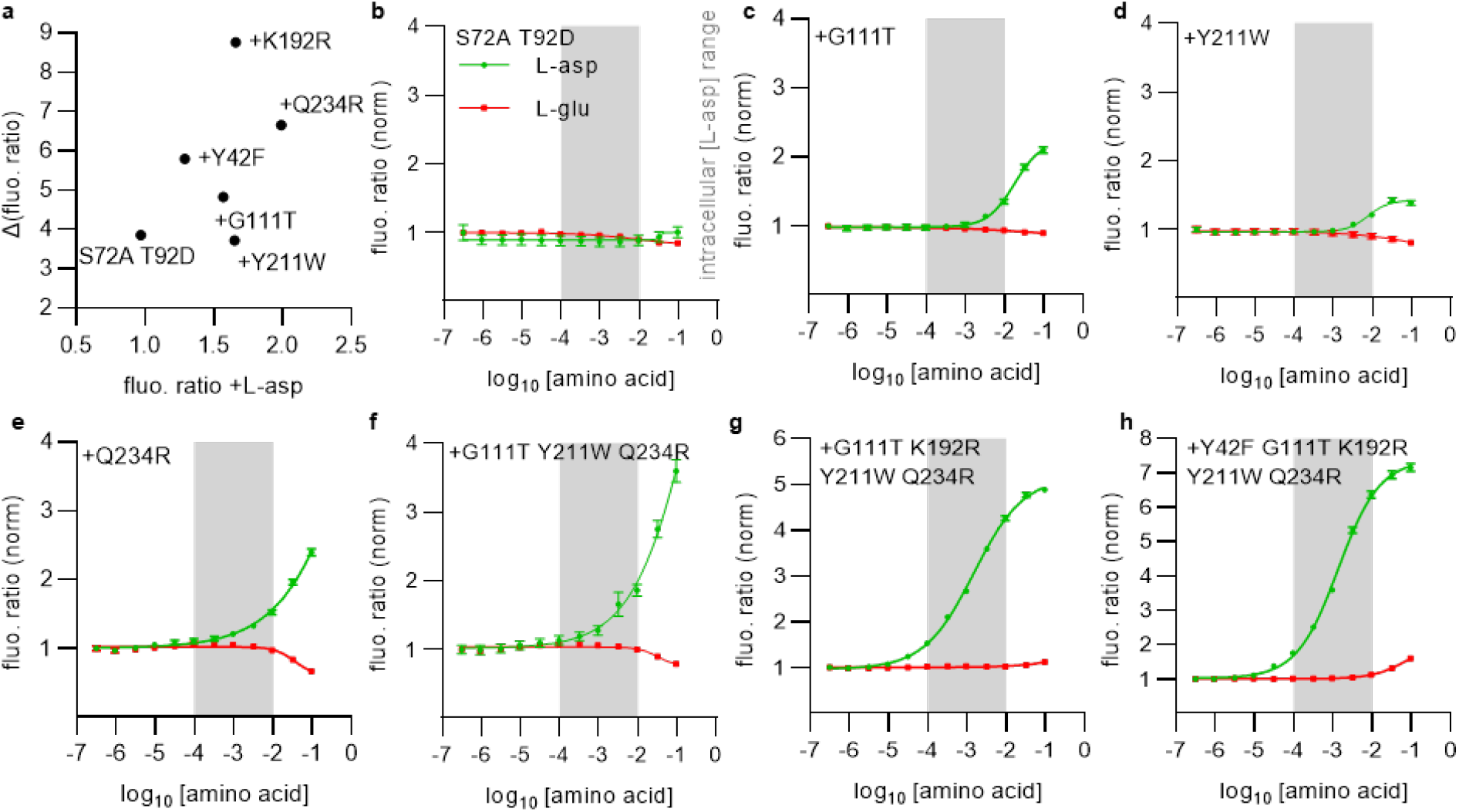
Stepwise iAspSnFR improvements by DMS mutations. **a)** DMS hits were validated in HEK293-JI cells displaying individual mutants of <u>cell-surface</u> SF-iGluSnFR-S72A T92D. All mutations increase absolute sensor fluo. in presence of aspartate (x-axis), and all except Y211W additionally improve the fluo. change (delta change in fluo. Ratio +/-aspartate, y-axis). Flow cytometry data, median of 40,000-55,000 cells. **b-h)** Lysate titrations of HEK293-JI cells stably expressing individual <u>cytoplasmic</u> mutants of SF-iGluSnFR-S72A T92D. **b)** SF-iGluSnFR-S72A T92D after manual screening, and prior to DMS. **c-f)** DMS#1 hits **g-h)** plus DMS#2 hits. **c-f)** To produce stable curve fits without reaching the upper curve plateau, the Hill slope was constrained to 1. **c-h)** Fit values in K_0.5_ / mM; fluo. ratio change / %: b) N/A c) 23, 140 d) 9, 50 e) 22, 160 f) 26, 310 g) 1.5, 420 h) 1.4, 640. Grey box: intracellular aspartate range.

**Supplementary Fig. 7.**
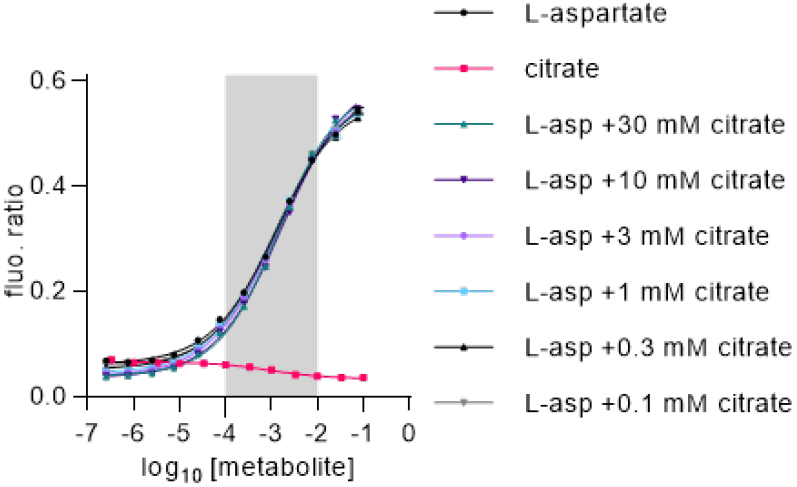
Aspartate titrations of purified iAspSnFR in presence of competing citrate. Grey box: intracellular aspartate range.

**Supplementary Fig. 8.**
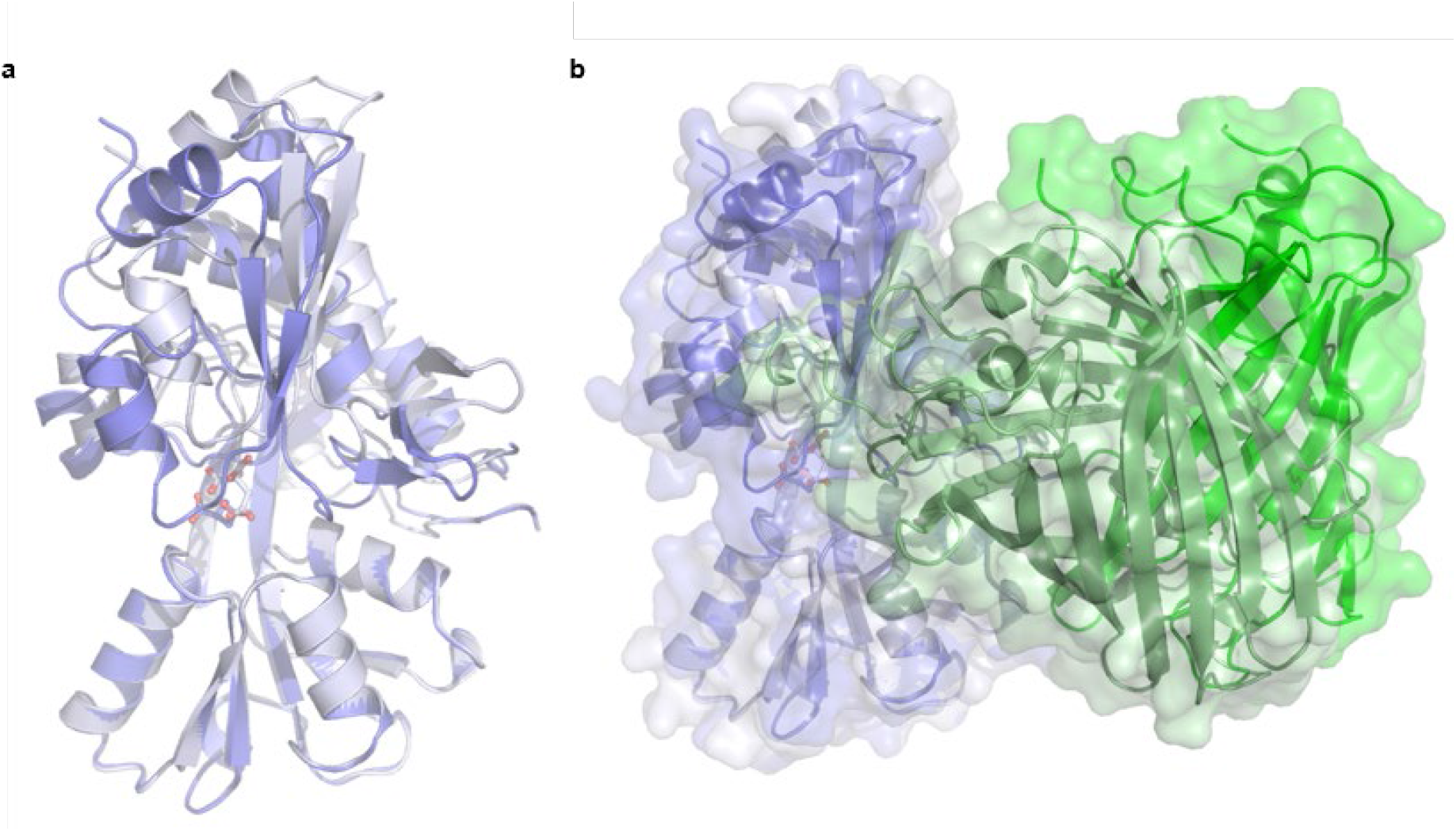
Crystal structures of SF-iGluSnFR-S72A. **a)** Shown is only the ligand binding domain. Overlay of aspartate (blue sticks)-bound (blue) and open-conformation with citrate (grey sticks) present in the binding pocket (grey) **b)** Overlay of the full sensor including the GFP domain (green), showing a large movement of the GFP domain in respect to the ligand binding domain upon aspartate binding.

**Supplementary Fig. 9.**
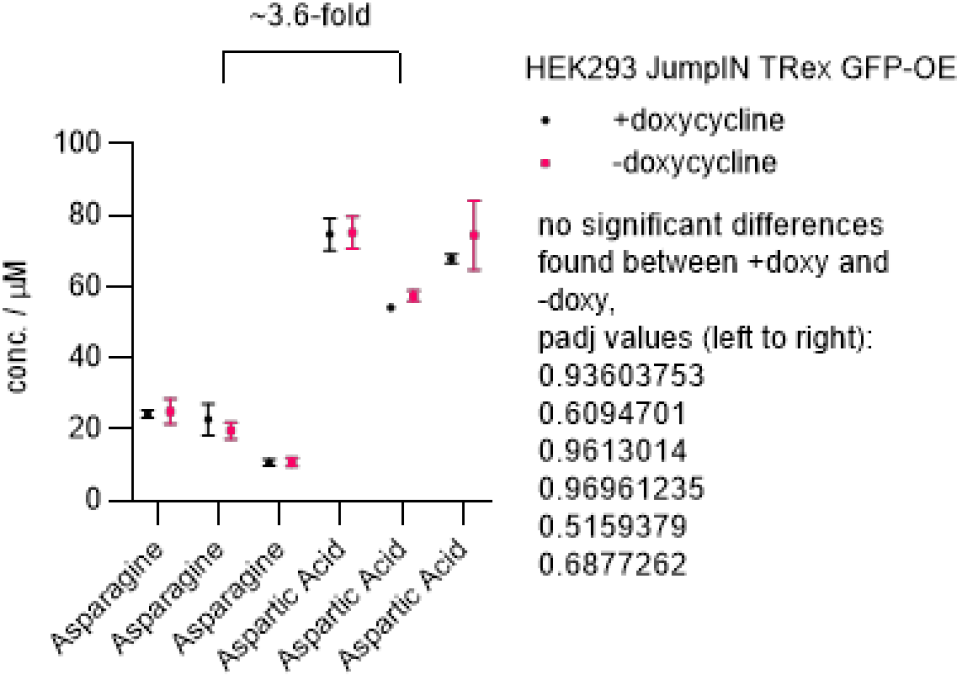
Metabolomics analysis (three biological replicates) of control HEK293-JI GFP-OE (doxycycline-inducible GFP-overexpression) shows a ∼3.6-fold ratio (average over all three biological replicates) of aspartate to asparagine concentrations. Moreover, no effect of doxycycline-induction of FP-expression on aspartate or asparagine levels was observed. Metabolomics datasets will be made publicly available at https://re-solute.eu/resources/datasets

**Supplementary Fig. 10.**
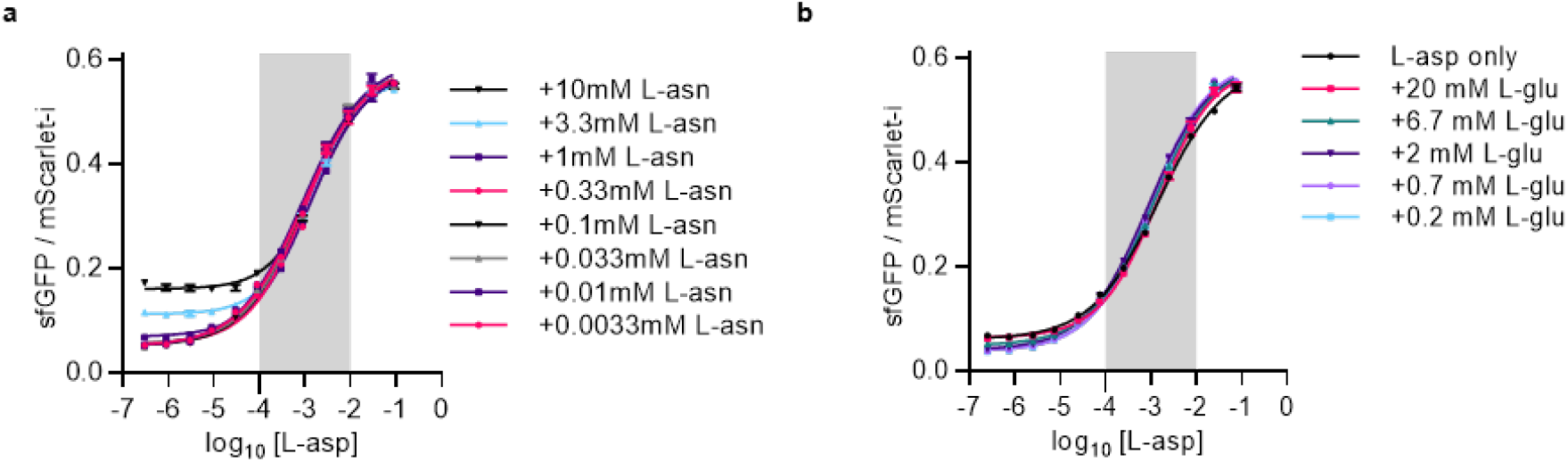
**a)** Aspartate titrations of purified iAspSnFR in presence of different concentrations of competing asparagine. **b)** Aspartate titrations of purified iAspSnFR in presence of different concentrations of competing glutamate. Grey box: intracellular aspartate range.

**Supplementary Fig. 11.**
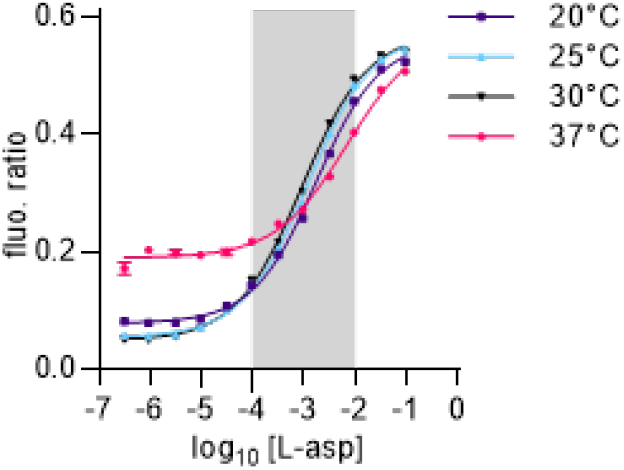
Purified iAspSnFR titrations at 20, 25, 30 and 37 °C.

**Supplementary Fig. 12.**
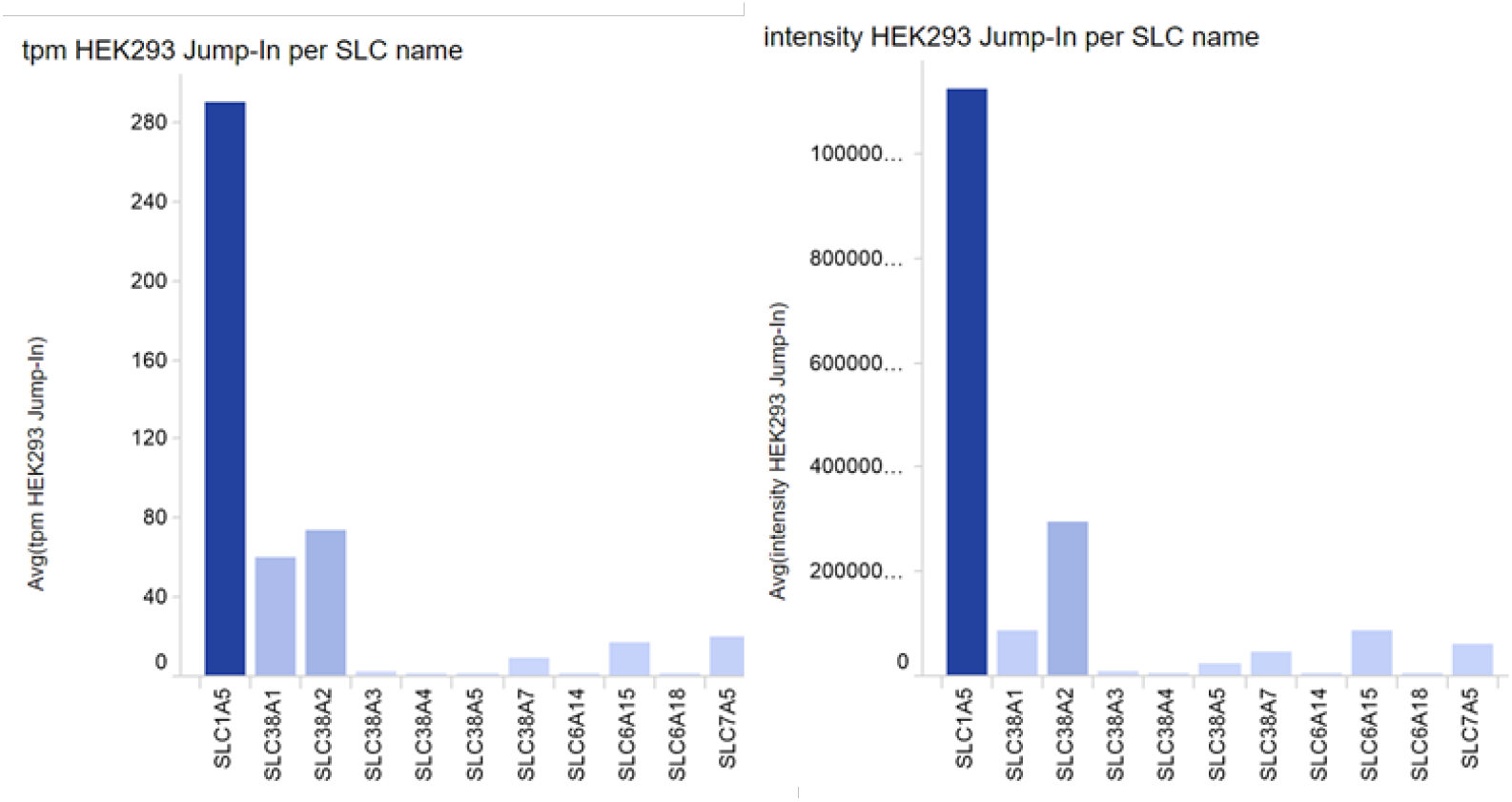
Abundancy of asparagine transporters in parental HEK293-JI cells by transcriptomics (left) and by proteomics (right). Datasets are publicly available for download at https://re-solute.eu/resources/datasets

## Notes

### Competing Interest Statement

The authors have declared no competing interest.

## References

1. Spinelli, J. B. & Haigis, M. C. The multifaceted contributions of mitochondria to cellular metabolism. Nat Cell Biol 20, 745–754 (2018).

2. Birsoy, K. et al.. An Essential Role of the Mitochondrial Electron Transport Chain in Cell Proliferation Is to Enable Aspartate Synthesis. Cell 162, 540–551 (2015).

3. Sullivan, L. B. et al. Supporting Aspartate Biosynthesis Is an Essential Function of Respiration in Proliferating Cells. Cell 162, 552–563 (2015).

4. Alkan, H. F. et al. Cytosolic Aspartate Availability Determines Cell Survival When Glutamine Is Limiting. Cell Metabolism 28, 706–720.e6 (2018).

5. Palmieri, L. et al. Citrin and aralar1 are Ca2+-stimulated aspartate/glutamate transporters in mitochondria. EMBO J 20, 5060–5069 (2001).

6. Borst, P. The malate–aspartate shuttle (Borst cycle): How it started and developed into a major metabolic pathway. IUBMB Life 72, 2241–2259 (2020).

7. Sullivan, L. B. et al. Aspartate is an endogenous metabolic limitation for tumour growth. Nature Cell Biology 20, 782 (2018).

8. Chen, W. W., Freinkman, E., Wang, T., Birsoy, K. & Sabatini, D. M. Absolute Quantification of Matrix Metabolites Reveals the Dynamics of Mitochondrial Metabolism. Cell 166, 1324–1337.e11 (2016).

9. Willis, R. C. & Furlong, C. E. Purification and properties of a periplasmic glutamate-aspartate binding protein from Escherichia coli K12 strain W3092. J. Biol. Chem. 250, 2574–2580 (1975).

10. De Lorimier, R. M. et al. Construction of a fluorescent biosensor family. Protein Science 11, 2655–2675 (2002).

11. Okumoto, S. et al. Detection of glutamate release from neurons by genetically encoded surface-displayed FRET nanosensors. Proceedings of the National Academy of Sciences 102, 8740–8745 (2005).

12. Deuschle, K. et al. Construction and optimization of a family of genetically encoded metabolite sensors by semirational protein engineering. Protein Science 14, 2304–2314 (2005).

13. Hires, S. A., Zhu, Y. & Tsien, R. Y. Optical measurement of synaptic glutamate spillover and reuptake by linker optimized glutamate-sensitive fluorescent reporters. PNAS 105, 4411–4416 (2008).

14. Marvin, J. S. et al. An optimized fluorescent probe for visualizing glutamate neurotransmission. Nature Methods 10, 162–170 (2013).

15. Helassa, N. et al. Ultrafast glutamate sensors resolve high-frequency release at Schaffer collateral synapses. PNAS 115, 5594–5599 (2018).

16. Marvin, J. S. et al. Stability, affinity, and chromatic variants of the glutamate sensor iGluSnFR. Nature Methods 15, 936–939 (2018).

17. Aggarwal, A. et al. Glutamate indicators with improved activation kinetics and localization for imaging synaptic transmission. 2022.02.13.480251 Preprint at https://doi.org/10.1101/2022.02.13.480251 (2022).

18. Fowler, D. M. & Fields, S. Deep mutational scanning: a new style of protein science. Nat Methods 11, 801–807 (2014).

19. Rebsamen, M. et al. Gain-of-function genetic screens in human cells identify SLC transporters overcoming environmental nutrient restrictions. Life Science Alliance 5, (2022).

20. Mendes, P., Oliver, S. G. & Kell, D. B. Fitting Transporter Activities to Cellular Drug Concentrations and Fluxes: Why the Bumblebee Can Fly. Trends Pharmacol Sci 36, 710–723 (2015).

21. Scalise, M., Pochini, L., Console, L., Losso, M. A. & Indiveri, C. The Human SLC1A5 (ASCT2) Amino Acid Transporter: From Function to Structure and Role in Cell Biology. Frontiers in Cell and Developmental Biology 6, (2018).

22. Zhang, H. et al. Comprehensive molecular and clinical characterization of SLC1A5 in human cancers. Pathology -Research and Practice 224, 153525 (2021).

23. Schalk, A. M., Nguyen, H.-A., Rigouin, C. & Lavie, A. Identification and Structural Analysis of an l-Asparaginase Enzyme from Guinea Pig with Putative Tumor Cell Killing Properties *. Journal of Biological Chemistry 289, 33175–33186 (2014).

24. Keshet, R., Szlosarek, P., Carracedo, A. & Erez, A. Rewiring urea cycle metabolism in cancer to support anabolism. Nat Rev Cancer 18, 634–645 (2018).

25. Vander Heiden, M. G. & DeBerardinis, R. J. Understanding the Intersections between Metabolism and Cancer Biology. Cell 168, 657–669 (2017).

26. Rabinovich, S. et al. Diversion of aspartate in ASS1-deficient tumours fosters de novo pyrimidine synthesis. Nature 527, 379–383 (2015).

27. Qi, L. et al. Aspartate availability limits hematopoietic stem cell function during hematopoietic regeneration. Cell Stem Cell 28, 1982–1999.e8 (2021).

28. Petrelli, F. et al. Mitochondrial pyruvate metabolism regulates the activation of quiescent adult neural stem cells. Sci Adv 9, eadd5220 (2023).

29. Tavoulari, S., Lacabanne, D., Thangaratnarajah, C. & Kunji, E. R. S. Pathogenic variants of the mitochondrial aspartate/glutamate carrier causing citrin deficiency. Trends in Endocrinology & Metabolism 33, 539–553 (2022).

30. Broeks, M. H., van Karnebeek, C. D. M., Wanders, R. J. A., Jans, J. J. M. & Verhoeven-Duif, N. M. Inborn disorders of the malate aspartate shuttle. Journal of Inherited Metabolic Disease 44, 792–808 (2021).

