## Supplementary Information for "Engineering of a biosensor for intracellular aspartate"

### Methods

#### Reagents, media and chemicals

Reagents, media and chemicals were obtained from different manufacturers and dissolved in suitable vehicles listed in **Table S1**. All compound and amino acid stocks were stored at -20°C, aliquoted to minimize freeze-thaw cycles, and freshly diluted prior usage.

**Table S1**: Reagents and Chemicals

| Chemical/Reagent | Manufacturer | Catalogue number | Notes |
|---|---|---|---|
| KOD Hot Start Master Mix | Sigma-Aldrich | 71842 | |
| BsmBI-v2 | NEB | R0739 | |
| BsaI-HF®v2 | NEB | R3733 | |
| Gibson assembly master mix | | | Home-made |
| QIAprep Spin Miniprep Kit | Qiagen | 27106 | |
| GeneJET Endo-Free Plasmid-Maxiprep-Kit | ThermoFisher | K0861 | |
| GenElute Mammalian Genomic DNA Miniprep Kit | Sigma-Aldrich | G1N350 | |
| HisPur™ Ni-NTA Superflow Agarose | ThermoScientific | 25217 | |
| Amicon® Ultra 4 mL Centrifugal Filters | Merck | UFC803024 (30 kDa) UFC805024 (50 kDa) | |
| TrypLE™ Express | Gibco | 12604013 | |
| DMEM high glucose +GlutaMAX™ | Gibco | 31966021 | |
| DMEM no glucose, phenol red-free | Gibco | 1443001 | |
| Sodium pyruvate (100X) | Gibco | 11360070 | |
| GlutaMAX™ Supplement (100x) | Gibco | 35050038 | |
| Fetal bovine serum (FBS, heat-inactivated) | Gibco | 10500064 | |
| Opti-MEM™ | Gibco | 31985047 | |
| HBSS+Ca+Mg | Corning | 21-023-CV | |
| PBS | Gibco | 10010023 | |
| Lipofectamine 3000 Transfection Reagent | Invitrogen | L3000001 | |
| D-glucose solution | Gibco | A2494001 | |
| Dialyzed Fetal Bovine Serum (dFBS), One-Shot | Gibco | A3382001 | |
| Doxycycline hyclate | Sigma-Aldrich | D9891 | 2 mg/mL in 70% EtOH, sterile filtered, single-use aliquoted |
| TFB-TBOA | Tocris | 2532 | DMSO, 25 mM |
| Rotenone | TRC Canada | R700580 | DMSO, 25 mM |
| Antimycin A | Sigma-Aldrich | A8674 | DMSO, 10mM |
| CB-839 | Medchemexpress | HY-12248 | DMSO, 10mM |
| L-amino acids | Sigma-Aldrich | LAA21 | PBS or HBSS; 100mM, pH=7.3-7.4 |
| D-aspartate | Carl Roth | 7873.2 | PBS or HBSS; 100mM, pH=7.3-7.4 |
| 96well plates, cell-grade, black, transparent F-bottom | BRANDplus | 735-2104 | For titrations using adherent live cells |
| 384well plates | Corning | 3573 | For titrations using purified proteins |
| 96well plates | PerkinElmer | 6055260 | For titrations using cleared lysate |
| CellLytic M | Sigma-Aldrich | C2978 | For mammalian cell lysis |
| COMPLETE(TM), MINI, EDTA-FREE PROTEASE Inhibitor cocktail | Roche | 11836170001 | 1 tablet per 10 mL CellLytic M buffer |

| | | | |
|---|---|---|---|
| Geneticin Selective Antibiotic 50mg/mL | Gibco | 10131027 | |
| Neon Transfection System 100uL | ThermoFisher | MPK10096 | |
| µ-Slide 18 Well Ibiditreat | Ibidi | 81816 | For microscopy experiments |
| µ-Slide VI 0.4 | Ibidi | 80604 | For perfusion experiments |

#### Plasmids and cloning

Oligonucleotides and synthetic genes, including the synthetic gene coding for SF-iGluSnFR-S72A, were obtained from multiple providers (IDT, Sigma-Aldrich, Genscript, Twist Bioscience, Eurofins). The KOD polymerase was used for most PCR amplifications. Cloning was performed either using TIIS restriction enzyme (NEB) cloning or Gibson assembly. Plasmid amplifications were performed in *E. coli* strains E. cloni 10G (Novagen), 5-alpha competent (NEB) or XL-1Blue (Agilent). Individual clones were picked, cultured and used for mini- or maxi-plasmid DNA extraction. Each plasmid preparation was verified by Sanger sequencing of the full insert. In some cases, *e.g.* plasmids used for key cell experiments, plasmids were additionally verified using whole-plasmid sequencing by Plasmidsaurus (Eugene, OR 97403). DNA concentration was measured using a Nanodrop 2000c spectrophotometer (Thermo Fisher). Plasmid DNA was aliquoted and stored in water at -20°C, avoiding excessive freeze-thaw cycles.

#### Protein expression and purification

Performed according to previously described procedures[1]. Very briefly, iAspSnFR was expressed from a pET51b(+) vector (Novagen) as N-terminal His-tag fusion in *E.coli* BL21 (DE3)-pLysS (Sigma-Aldrich). Ni-NTA affinity purification was followed by TEV-cleavage of the His tag, producing the same protein sequence as used in mammalian cell experiments except that the *N*-terminal methionine is replaced by a glycine. The protein was further purified by Ni-NTA purification to scavenge uncleaved protein and the flow through was subjected to size exclusion chromatography (SEC). Protein purity was assessed by protein gel and SEC chromatography profile, protein species identity was confirmed using mass spectrometry. The pure protein was used for crystallization, photophysical characterization and titrations. Protein concentration was determined by measuring absorbance at 280 nm using a Nanodrop 2000c spectrophotometer (Thermo Fisher).

Purified iAspSnFR sequence after TEV cleavage:

GVSKGEAVIKEFMRFKVHMEGSMNGHEFEIEGEGEGRPYEGTQTAKLKVTKGGPLPFSWDILSPQFMYGSRA
FIKHPADIPDYYKQSFPEGFKWERVMNFEDGGAVTVTQDTSLEDGTLIYKVKLRGTNFPPDGPVMQKKTMGW
EASTERLYPEDGVLKGDIKMALRLKDGGRYLADFKTTYKAKKPVQMPGAYNVDRKLDITSHNEDYTVVEQYER
SEGRHSTGGMDELYKGS**AAGSTLDKIAKNGVIVVGHRESSVPFSYYDNQQKVVGFSQDYSNAIVEAVKKKLN**
**KPDLQVKLIPITAQNRIPLLQNGTFDFECGSTDNNVERQKQAAFSDTIFVVTTRLLTKKGGDIKDFANLKDKAV**
**VVTSGTTSEVLLNKLNEEQKMNMRIISAKDHGDSFRTLESGRAVAFMMDDVLLAGERARAKKPDNWEIVGK**
**PQSQEAWGCMLRKDDPQFKKLMDDTIAQVRTSGEAEKWFDKWFKNPILV**SHNVYITADKQKNGIKANFKIRH
NVEDGSVQLADHYQQNTPIGDGPVLLPDNHYLSTQSVLSKDPNEKRDHMVLLEFVTAAGITLGMDELYKGGTG
GSMSKGEELFTGVVPILVELDGDVNGHKFSVRGEGEGDATNGKLTLKFICTTGKLPVPWPTLVTTLTYGVQCF

SRYPDHMKQHDFFKSAMPEGYVQERTISFKDDGTYKTRAEVKFEGDTLVNRIELKGIDFKEDGNILGHKLEYNF
N**NPLNMNFELSDEMKALFKEPNDKALK**

mScarlet-I, cp-sfGFP, **gltI binding protein**

Purified iAspSnFR lacking mScarlet-I, construct used for protein crystallization and X-ray crystallography:

G**AAGSTLDKIAKNGVIVVGHRESSVPFSYYDNQQKVVGFSQDYSNAIVEAVKKKLNKPDLQVKLIPITAQNRIP
LLQNGTFDFECGSTDNNVERQKQAAFSDTIFVVTTRLLTKKGGDIKDFANLKDKAVVVTSGTTSEVLLNKLNE
EQKMNMRIISAKDHGDSFRTLESGRAVAFMMDDVLLAGERARAKKPDNWEIVGKPQSQEAWGCMLRKDDP
QFKKLMDDTIAQVRTSGEAEKWFDKWFKNPILV**SHNVYITADKQKNGIKANFKIRHNVEDGSVQLADHYQQNT
PIGDGPVLLPDNHYLSTQSVLSKDPNEKRDHMVLLEFVTAAGITLGMDELYKGGTGGSMSKGEELFTGVVPILV
ELDGDVNGHKFSVRGEGEGDATNGKLTLKFICTTGKLPVPWPTLVTTLTYGVQCFSRYPDHMKQHDFFKSAM
PEGYVQERTISFKDDGTYKTRAEVKFEGDTLVNRIELKGIDFKEDGNILGHKLEYNFN**NPLNMNFELSDEMKAL
FKEPNDKALK**

cp-sfGFP, **gltI binding protein**

**Mass spectrometry**

Samples were fractionated using a Shimadzu Nexera HPLC. The protein sample concentrations was adjusted to 10 μM. The resulting solution was injected on a Phenomenex, Aeris C4 (2.1x100 mm, 3.6 μ, widepore) column operated at 50 C using an injection volume of 5 μL. Target protein elution was facilitated using a total flow rate of 400 μL/min of mobile phase A, water with 0.1 % formic acid, and phase B, acetonitrile with 0.1 % formic acid (LCMS grade). The ratio of the two solvents was adjusted according to the following gradient profile: 0 min 10% B, 1min 10% B, 6 min 98% B, 7.5 min 98 % B, 7.6 min 10% B and 8.6 min 10% B. A sodium formate solution was injected within the time range of the re-equilibration plateau to allow for a sodium formate cluster based internal calibration of the acquired MS spectra.

The HPLC eluates were analyzed with a Bruker maXis II ETD high resolution mass spectrometer equipped with its high flow source set to the following parameters: End plate offset 500 V, capillary voltage 4500 V, nebulizer pressure 40 psi, dry gas flow 10 L/min and a try temperature of 250 C. Spectra were acquired in a mass range from 400 to 4000 m/z with a spectra rate of 1.0 Hz. The hardware was controlled with Brukers Hystar 5.1 and otofControl 5.2 software. The subsequent data analysis was done with Brukers DataAnalysis 5.3 software.

After internal calibration, an average mass spectrum for the time range 3 to 4.3 min was created. The resulting data was deconvoluted using the Maximum Entropy (MaxEnt) algorithm with the following settings: Low mass 10000, high mass 185000, data point spacing: Auto, resolving power 8000 and resolution high. Peakfinding in deconvoluted spectra was achieved with the SumPeak algorithm.

Purified iAspSnFR sequence after TEV cleavage was analyzed by mass spectrometry: Expected mass 84,436 Da – 42 Da = 84,394 Da (expected mass shift due to chromophore maturation[2]: 2x21=42 Da). Observed mass: 84,394 Da

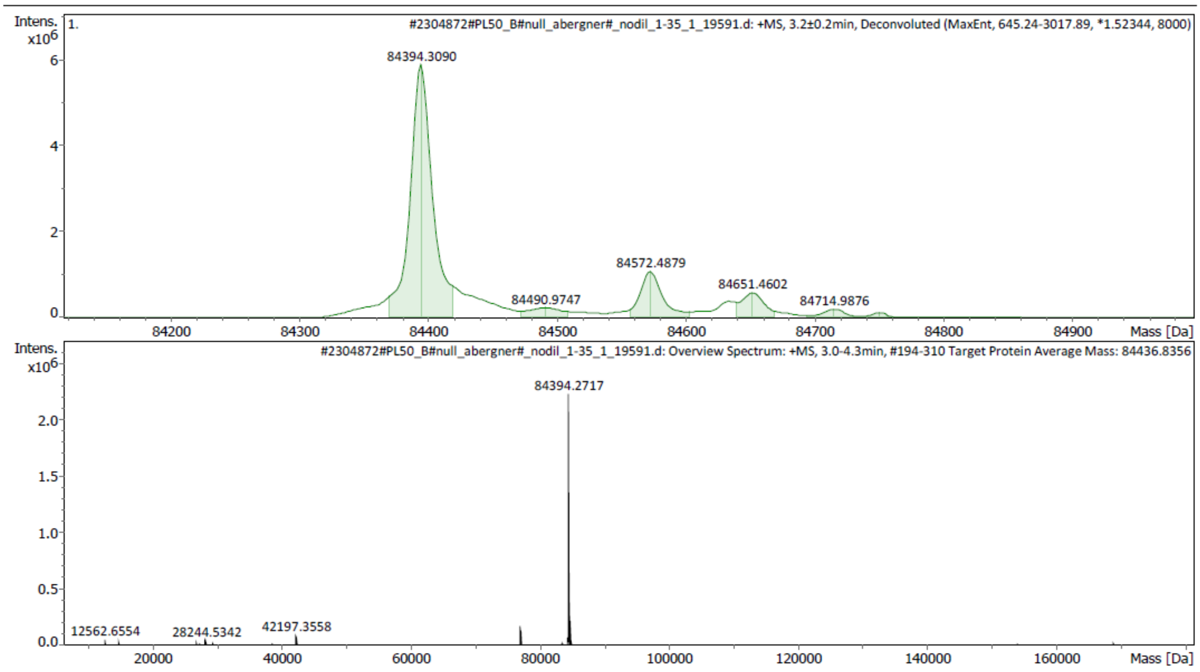

**Supplementary Fig. 13** | Mass spectrogram for iAspSnFR.

#### *In vitro* titrations using purified iAspSnFR

Measurements were performed in black, flat bottom, polystyrene 384-well plates in HBSS (except the pH titration) with final protein concentration at 100 nM on a Multimode Spark 20M microplate reader (Tecan) in top reading mode. Temperature was 20°C unless stated differently.

The pH titrations (pH = 5, 6, 7, 8, 9, 10) were performed in a MOPS/citrate/borate buffer (in mM: 30 MOPS, 30 citric acid, 30 sodium tetraborate decahydrate, 100 NaCl). L-aspartate was added as solid to a concentration of 10 mM, before setting the pH with NaOH or HCl. To control osmolarity, D-aspartate was added at equimolar concentration to the “buffer-only” sample.

**Table S2**: Microplate reader settings

| Parameter | Value |
|---|---|
| **sfGFP** | |
| Excitation / nm | 485/20 (filter) |
| Mirror | Dichroic 510 |
| Emission / nm | 510/10 (filter) |
| **mScarlet-I** | |
| Excitation / nm | 535/25 (filter) |
| Mirror | Dichroic 560 |
| Emission / nm | 595/35 (filter) |

**Photophysical characterization**

Quantum yields were determined using a Quantaurus QY (Hamamatsu). iAspSnFR+Asp was measured 250 nM and apo-iAspSnFR at 2 μM (in HBSS). QY was measured at 488nm excitation. Reported as mean±SD of four replicates.

For measurement of sfGFP extinction coefficients, significant overlap of mScarlet-I and sfGFP absorbance was observed, preventing accurate measurement of absorbance at 494 nm (A494). Therefore, for accurate measurement, protein was produced and purified lacking the *N*-terminal mScarlet-I domain and used for measurement of A494. Extinction coefficients were determined by measuring absorbance at the indicated wavelengths using a Nanodrop 2000c spectrophotometer.

Excitation and emission spectra were measured at Multimode Spark 20M microplate reader (Tecan) in top reading equipped with a monochromator. Spectra were normalized to the peak excitation or emission.

**Protein crystallization**

Crystallization was performed at 20°C using the vapor-diffusion method. SF-iGluSnFR-S72A and iAspSnFR were used at a concentration of 12.0 mg/ml in 50 mM HEPES pH 7.3, 50 mM sodium chloride. Crystals of SF-iGluSnFR-S72A were grown by mixing protein solution and a reservoir solution containing 1.5 M tri-sodium citrate pH 6.5 at a 2:1 ratio. Crystals of SF-iGluSnFR-S72A in complex with L-aspartate were obtained by mixing equal volumes of protein solution supplemented with 20 mM L-aspartate and precipitant solution composed of 0.2 M lithium acetate, 21% (m/v) PEG 3350. Crystals of iAspSnFR in complex with L-aspartate were grown by mixing equal volumes of protein solution supplemented with 40 mM L-aspartate and precipitant solution containing 0.2 M magnesium acetate, 18% (m/v) PEG 3350. Crystals of SF-iGluSnFR-S72A were briefly washed in cryoprotectant solution consisting of the reservoir solution supplemented with 10% (m/v) glucose and 10% (m/v) sucrose before flash-cooling in liquid nitrogen, whereas crystals of SF-iGluSnFR-S72A in complex with L-aspartate were washed in the reservoir solution with PEG 3350 added to a final concentration of 40% (m/v). Crystals of iAspSnFR in complex with L-aspartate were rinsed in the reservoir solution supplemented with 20% (v/v) glycerol, prior to flash-cooling in liquid nitrogen.

**X-ray diffraction data collection and structure determination**

Single crystal X-ray diffraction data were collected at 100 K on the X10SA beamline at the SLS (PSI, Villigen, Switzerland). All data were processed with XDS[3]. The structure of SF-iGluSnFR-S72A was determined by molecular replacement (MR) with Phaser[4] using aspartate/glutamate binding protein (DEBP) from *Shigella flexneri* (PDB code 2VHA) and GFP (PDB code 6EFR) coordinates as a search models. The structures of SF-iGluSnFR-S72A and iAspSnFR in complex with L-aspartate were determined using the SF-iGluSnFR-S72A model. The final models were optimized in iterative cycles of manual rebuilding using Coot[5] and refinement using Refmac5[6] and phenix.refine[7]. Data collection and refinement statistics are summarized in **Table S3**, model quality was validated with MolProbity[8] as implemented in PHENIX.

Atomic coordinates and structure factors have been deposited in the Protein Data Bank under accession codes: 8OVN (SF-iGluSnFR-S72A), 8OVO (SF-iGluSnFR-S72A:L-Asp), 8OVP (iAspSnFR:L-Asp).

**Table S3**: Data collection and refinement statistics. Values in parentheses are for the highest resolution shell.

| | SF-iGluSnFR-S72A<br>PDB code: 8OVN | SF-iGluSnFR-S72A:L-Asp<br>PDB code: 8OVO | iAspSnFR:L-Asp<br>PDB code: 8OVP |
|---|---|---|---|
| **Data collection** | | | |
| Space group | $P3_121$ | $P2_12_12_1$ | $P12_11$ |
| Unit-cell parameters | | | |
| *a, b, c* (Å) | 106.14, 106.14, 108.14 | 65.76, 77.48, 196.84 | 67.31, 105.88, 77.28 |
| $\alpha, \beta, \gamma$ (°) | 90.00, 90.00, 120.00 | 90.00, 90.00, 90.00 | 90.00, 92.45, 90.00 |
| Radiation source | PXII-X10SA, SLS | PXII-X10SA, SLS | PXII-X10SA, SLS |
| Wavelength (Å) | 1.00007 | 1.00008 | 0.99998 |
| Temperature (K) | 100 | 100 | 100 |
| Resolution range (Å) | 50-2.60 (2.70-2.60) | 50-1.70 (1.80-1.70) | 50-1.70 (1.80-1.70) |
| No. of observed reflections | 185962 (20280) | 830713 (132945) | 405064 (64501) |
| No. of unique reflections | 22091 (2344) | 111306 (17344) | 116407 (18188) |
| Multiplicity | 8.4 (8.7) | 7.5 (7.7) | 6.3 (6.4) |
| Completeness (%) | 99.9 (100.0) | 99.9 (100.0) | 98.1 (97.5) |
| $R_{merge}$ (%) | 8.3 (71.0) | 3.6 (60.7) | 3.9 (54.8) |
| $\langle I/\sigma(I) \rangle$ | 19.3 (3.2) | 27.7 (3.2) | 16.7 (2.3) |
| $CC_{1/2}$ (%)[#] | 99.9 (85.8) | 100.0 (87.3) | 99.9 (77.6) |
| **Refinement** | | | |
| Molecules per a.u. | 1 | 2 | 2 |
| No. of reflections | 22086 | 111305 | 116402 |
| No. of reflections in test set | 1105 | 5566 | 5821 |
| Resolution range (Å) | 46.60-2.60 | 48.59-1.70 | 38.60-1.70 |
| No. of non-hydrogen atoms | | | |
| Protein | 3965 | 8013 | 8006 |
| Ligand/ion | 35 | 44 | 55 |
| Water | 19 | 731 | 684 |
| Total | 4019 | 8788 | 8745 |
| *R* (%) | 19.96 | 19.03 | 17.14 |
| $R_{free}$ (%) | 25.28 | 22.00 | 20.16 |
| RMS deviations from ideal | | | |
| bonds (Å) | 0.002 | 0.007 | 0.009 |
| angles (°) | 0.521 | 0.898 | 1.009 |
| *B*-factors (Å$^2$) | | | |

| | | | |
|---|---|---|---|
| Protein | 49.36 | 30.96 | 29.72 |
| Ligand/ion | 52.23 | 25.20 | 25.57 |
| Water | 43.12 | 35.61 | 36.80 |
| Average | 49.36 | 31.32 | 30.25 |
| Wilson B ( Å$^2$) | 51.42 | 28.72 | 26.61 |
| Ramachandran statistics (%) | | | |
| favored regions | 97.8 | 97.9 | 98.3 |
| allowed regions | 2.2 | 2.1 | 1.7 |
| disallowed regions | 0 | 0 | 0 |
| Clashscore | 3.02 | 2.12 | 2.73 |

#as implemented in XDS[9].

**Cell culture**

Full medium refers to 10% FCS/DMEM high-glucose, pyruvate, glutamax, phenolred (Gibco #31966021), used for standard cell culture.

Reduced base medium refers to 10% dialyzed FCS/DMEM (Gibco #1443001). It was freshly supplemented as indicated (e.g. pyruvate/glucose/glutamax) and used as experimental medium. The glutamax supplement at 1x was used instead of glutamine for all experiments.

Starvation medium refers to 10% dialyzed FCS/DMEM (Gibco #1443001).

All media were prepared from reagents listed in **Table S1**, were sterile filtered prior usage (Nalgene #564-0020), and cultured without Pen/Strep. Cells were cultured at 37 °C and 5% $CO_2$ in a humified incubator. Cells were regularly checked for mycoplasma infection by PCR and always tested negative.

**Table S4**: Cell lines

| Chemical/Reagent | Source | Identifier | Notes |
|---|---|---|---|
| HEK293 Jump In™ T-Rex™ (abbreviated as HEK293-JI) | RESOLUTE / Thermo Fisher | CE0002-U | Datasheet ThermoFisher: https://www.thermofisher.com/order/catalog/product/A15008?SID=srch-srp-A15008 |
| HEK293 Jump In™ T-Rex™ SLC1A3-WTOE | RESOLUTE | CE029W-F | https://re-solute.eu/files/cellline_reports/CE029W-F.pdf |

**Stable cell line generation**

For stable cell line generation, plasmids encoding the R4 integrase and transgene was electroporated using the Neon Transfection System (Invitrogen) with 100 μL tips according to the manufacturer's instructions. Per individual electroporation, 2 Mio HEK293-JI cells, 3 μg R4 integrase and 5 μg transgene was used. For the DMS library, a total of 5x2=10 Mio (DMS#1) or 10x2=20 Mio (DMS#2) cells were used.

Two days after electroporation, HEK293-JI cells were selected with 2 mg/mL Geneticin in full medium until cell death ceased (~7-10 days). After the initial selection, cells were cultured without Geneticin as no significant loss of transgene was observed.

**Fluorescence-activated cell sorting (FACS)**

HEK293-JI cells with individual sensor variants or sensor libraries in the R4 site were induced with 1 μg/mL doxycycline over 1-2 days. Cells (one confluent T75 flask per sample) were washed with PBS, briefly trypsinized, and taken into suspension using reduced base medium. Cells were collected by centrifugation (800 rpm), supernatant was removed, and cells were resuspended in 10% dialyzed FCS/PBS before filtration through a cell strainer. Cells were sorted using a BD FACS Melody instrument (sfGFP/FITC channel: Ex 488 nm, Em 527/32; mScarlet-I/mCherry channel: Ex 561, Em 613/18). Singlets were isolated by gating SSC-A vs FSC-A and then by FSC-W vs FSC-H. mScarlet-I positive cells were gated for subsequent analysis and cell sorts.

If aspartate or glutamate was added to increase fluo. ratio of iAspSnFR-variant displaying cells, it was added from a 100mM PBS stock to a saturating nominal concentration of 33.3 mM.

Prior to every experimental day, the HEK293-JI cell line expressing the “wild type” SF-iGluSnFR-S72A T92D cell-surface screening construct was run in presence or absence of aspartate as control.

For enrichments by cell sorting, gates were drawn in the sfGFP vs mScarlet-I plot and manually adjusted so that the appropriate (and approximate) percentage of cells ended up in the collection tube (e.g. top 1% sfGFP/mScarlet-I ratio). Enriched/sorted cells were taken back into culture and grown back to a confluent T75 flask before performing the next enrichment step as necessary.

Data analysis was performed in FlowJo 10. Fluo. ratio was derived by dividing GFP:A by mScarlet:A, and depicted as histogram with modal y-axis. Gating strategy: 1) live cells in FSC:A vs SSC:A 2) mSc positive cells in SSC:A vs mScarlet:A.

Example gating for control (HEK293 JumpIN TRex cell line expressing the “wild type” cell-surface screening construct):

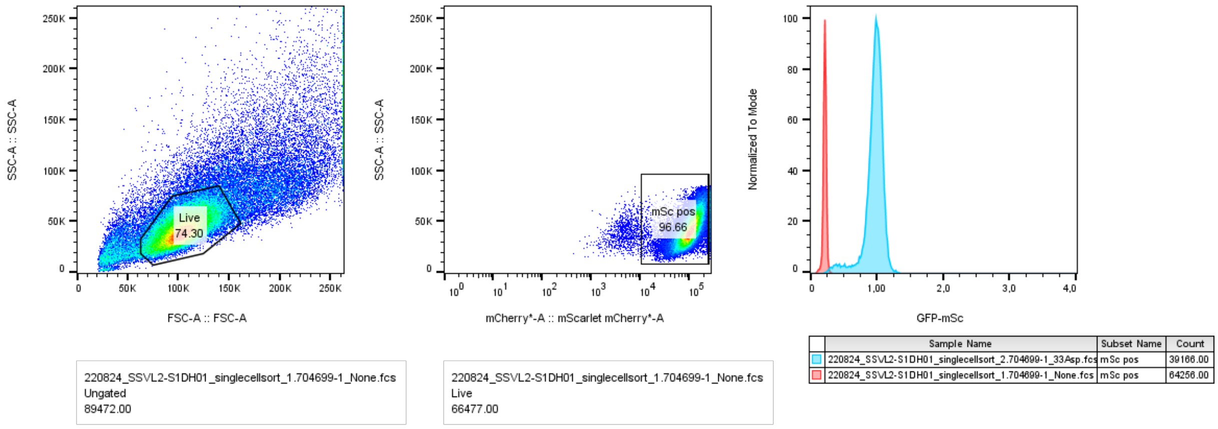

**Supplementary Fig. 14 |** FACS example gating for control (HEK293-JI cell line expressing the "wild type" cell-surface screening construct).

#### Lysate titrations of HEK293-JI stably expressing sensor variants

Per sample, one 10 cm$^2$ dish at 60-90% confluency was used and lysate for titrations was always prepared freshly. Medium was removed, cells were washed with 4°C cold PBS, then 300-500 µL ice-cold lysis buffer (CellLytic M supplemented with protease inhibitor, see Table S1) was added and cells were scraped off. Lysate was transferred to 1.5 mL Eppendorf tubes, briefly vortexed, and kept on ice. Samples were centrifuged at >20,000 rpm and 4°C for 20 min. The cleared lysate was transferred to a fresh 1.5 mL Eppendorf tube without perturbing the debris pellet and directly used for subsequent titrations. Titrations were performed in 96-well plates using 5 µL/well of cleared protein lysate plus 100 µL/well total volume (HBSS buffered). Measurements were performed on Multimode Spark 20M microplate reader (Tecan) in top reading mode. Temperature was 20°C unless stated differently. Filter settings are the same as detailed in Table S2.

#### Genomic DNA extraction and PCR amplification prior to Illumina sequencing

Ca. 2 Mio cells were harvested by centrifugation (800 rpm, 10 min) and media was removed. Extraction of gDNA was performed using the GenElute Mammalian Genomic DNA Miniprep Kit (Sigma-Aldrich #G1N350) according to the manufacturer's instructions. gDNA was quantified and then amplified in two PCR steps (primer sequences in **Table S5**):

Step1: Amplification of the full biosensor binding domain with Pr697+Pr653.

Step2: Using the purified PCR product from step 1 as template, PCR amplification for Amplicon sequencing was performed using NGS-quality primers (Pr735-750) with 10-mer sample barcodes on the forward primer. This yields 4 overlapping amplicons per sample between nucleobases 17-259, 161-431, 376-623, 562-823.

Illumina adapter ligation and Illumina sequencing were performed by Eurofins (Ebersberg, Germany) in NovaSeq 6000 S4 PE150 XP sequence mode.

**Table S5**: Primers for Illumina sample preparation

| Primer | Sequence |
|---|---|
| Pr697_IgK_F | GTACTGCTGCTCTGGGTTCC |
| Pr653_L1_rev | CCTCCACGTTGTGGCGGATCTTG |
| Pr735_SSVL_17F_A | TTCACGGAAGCAGGTTCCACTGGTGACCG |
| Pr736_SSVL_17F_B | ATCGACGGCTCAGGTTCCACTGGTGACCG |
| Pr737_SSVL_17F_C | GTAAGGCTCCCAGGTTCCACTGGTGACCG |
| Pr738_SSVL_259R | GAATACGGTTTTGCGCGGTAAT |
| Pr739_SSVL_161F_A | CCTCGAATGGAGGATTACTCCAACGCCATTGT |
| Pr740_SSVL_161F_B | TCATGGAATCAGGATTACTCCAACGCCATTGT |
| Pr741_SSVL_161F_C | ATAAGAGGTCAGGATTACTCCAACGCCATTGT |
| Pr742_SSVL_431R | ACTACGGCTTTGTCTTTCAGGT |
| Pr743_SSVL_376F_A | GCTGACCTGAACCAAAAAGGGTGGCGATATCA |
| Pr744_SSVL_376F_B | CGACGTAGTCACCAAAAAGGGTGGCGATATCA |
| Pr745_SSVL_376F_C | TCAATGATCGACCAAAAAGGGTGGCGATATCA |
| Pr746_SSVL_623R | TTGTCTGGTTTCTTCGCTTTCG |
| Pr747_SSVL_562F_A | AGTCTCGGCAGCCTTTATGATGGATGACGTGC |
| Pr748_SSVL_562F_B | GATATAGCTCGCCTTTATGATGGATGACGTGC |
| Pr749_SSVL_562F_C | CGTCCGACTTGCCTTTATGATGGATGACGTGC |
| Pr750_SSVL_837R | CTTGATGCCGTTCTTCTGCTTG |

**Illumina sequencing analysis**

DMS sequencing analysis proceeded using a combination of command line tools and python packages. Briefly, Je demultiplex[10] was used for barcode sample separation, followed by BBMerge[11] to merge paired-end reads. BBTools/reformat.sh was used to reverse-complement reads where the barcode was found at the 3' end (since adapter ligation produces a random orientation of sample barcode in respect to indexing adapters). The reverse-complemented and the correctly-oriented reads were concatenated to obtain one fastq file per amplicon. Next, cutadapt[12] was called to remove primer sequences. The processed reads were analyzed in a custom python script, using an adaptation of mutagenesis-visualization[13], returning variant frequencies (defined as variant reads/all reads).

**Cell preparation for steady state microscopy (Fig. 3a, b, e, g, i)**

For steady-state microscopy experiments, the base medium was 10% dialyzed FCS/DMEM (Gibco #1443001), supplemented with 1X pyruvate, 1X glutamax or 4.5 g/L glucose as indicated. Instead of glutamine, all experiments were performed using the glutamax (200 mM L-alanyl-L-glutamine-dipeptide) supplement.

Cells were seeded at day 0 on μ-Slide 18 Well Ibiditreat microscopy dishes at 10,000-15,000/well in 100 μL full medium (referring to 10% FCS/DMEM Gibco #31966021) containing

1 μg/mL doxycycline. If the experiment requires transfection, at day 1 medium was removed by suction and the transfection mix was added (Lipofectamine 3000, 100 ng DNA, 0.2 μL P3000, 0.15 μL R3000, 10+10 μL OptiMEM) in 100 μL full medium with doxycycline reduced to 0.1 μg/mL. If the experiment doesn't require transfection, medium was removed by suction and replaced by 100 μL full medium with doxycycline reduced to 0.1 μg/mL.

On day 2, for treatments, medium was completely removed by suction, then compound-containing base medium (referring to 10% dialyzed FCS/DMEM Gibco #1443001) was added. Cells were incubated for 3 hrs (for nutrient deprivation, an increase to overnight incubation only slightly further reduced fluo. ratio) at 37 °C and 5% $CO_2$, before sample transferal to the microscope at 30 °C and 5% $CO_2$, allowing 15 mins for the dish to equilibrate. For every well, the bottom of the cell layer was manually found by adjusting z, then one z-stack (bottom to top, 11 slices over 10 μm) was recorded.

**Timelapse microscopy (Fig. 3c, d)**

Timelapse microscopy of HEK-SLC1A3 was performed in saline (in mM: 146 NaCl, 5 KCl, 1.2 $MgCl_2$, 2.5 $CaCl_2$, HEPES 5, pH 7.4) at 30 °C without $CO_2$. The cell bottom was manually found by adjusting z, and one z-stack (bottom to top, 11 slices over 10 μm) was recorded every 5 minutes. At the indicated timepoint, aspartate was manually added to the well with an Eppendorf pipette.

**Microscopy settings**

Confocal microscopy was performed on a Stellaris 5 inverted microscope (Leica) equipped with a white line laser and hybrid photodetectors at 30 °C in 5% $CO_2$ atmosphere in a humidified chamber. An HC PL APO CS2 63x/1.40 oil immersion objective was used at to image a z-stack (11 slices, 10 μm) at 1024x1024 pixel (185x185 μm) resolution (400 Hz scan speed, 95.6 μm pinhole) and with 16-bit depth. Channels were set as follows in line sequential mode: sfGFP: Ex 489 nm, Em 494-550 nm; mScarlet-I: Ex 569 nm, Em 580-750 nm. Laserpower was adjusted to avoid pixel saturation and ranged between 0.25-2%. For HEK293-JI-iAspSnFR cells, 1% was used for both excitation lines. Laserpower for both excitation lines was always adjusted simultaneously (i.e. 0.5% 489 nm and 0.5% 569 nm; 1% 489 nm and 1% 569 nm and so on), the fluo. ratio remained constant in this range (0.25-2%).

**Image processing**

Batch processing of microscopy images was performed using a pipeline in FIJI/JIPipe[14]. Image z-stacks were imported and an average intensity projection was performed, then fluo. channels were split. The mScarlet-I channel was used to perform cell segmentation by calling cellpose[15] using the cyto2torch_0 model with average diameter set at 130, and minimum size 10. The returned ROIs were used to extract the mean/min/max pixel values per cell on the individual channels, which were exported as csv files. This yields one csv file per image and channel. Data wrangling was performed in a custom JupyterLab notebook. The csv files were imported and combined as a list of pandas dataframes (one dataframe per well). Then, the fluo. ratio was calculated by dividing Mean_GFP by Mean_mSc. Cells were filtered for pixel values as follows: Mean_mSc > 1000 and Max_GFP < 65000 Max_mSc < 65000. This removes from the analysis non-expressing cells, and cells with saturated pixels that would skew the fluo. ratio. Lastly, the median of fluo. ratio, together with standard error (STD) and cell number (N), of each well was exported as Excel table. Those files were then used for visualization and statistical testing using Graphpad Prism 9.

**Flow cytometry (Fig. 3f, h)**

HEK293-JI SLC1A3-WTOE or HEK293-JI iAspSnFR cells were seeded and induced in full medium with 1 μg/mL doxycycline on day 0, and transiently transfected with pcDNA3.1/iAspSnFR or pSF-EF1a/EBFP-gpASNase1 on day 1 using Lipofectamine 3000 and changing medium to reduced base medium supplemented with glutamax, pyruvate, glucose and 0.1 μg/mL doxycycline. On day 2, cells were starved in reduced base medium for 3 hrs. Then, medium was removed, cells were washed with PBS, briefly trypsinized, and suspended in reduced base medium. Cells were collected by centrifugation, medium was removed, and resuspended in 10% dialyzed FCS/PBS before filtering through a cell strainer. The cell suspension was then dispensed into 96-well U-bottom plates suitable for the flow cytometer's HTS autosampler module. Compounds were added directly to the 96-well plate and incubated for ~30 min at room temperature (22°C), before measurement was initialized at a BD Fortessa X-20 flow cytometer. Optical configuration: EBFP/BV421 Ex 405 nm Em 450/50 nm, sfGFP/FITC Ex 488 nm Em 530/30nm, mScarlet-I/PE-Texas Red Ex 561 nm, Em 610/20 nm. 80-100 μL were injected per sample and all events were recorded and stored for subsequent analysis in FlowJo 10. Gating for the SLC1A3 was performed as described for FACS, fluo. ratio was derived by dividing FITC-A/PE-Texas Red-A. For the EBFP-gpASNase1 experiment, cells were additionally gated in the BV421-A channel to separate the EBFP-gpASNase1 -positive from negative cells. Transfection efficiency was tuned to 20-30%, so that both cell populations can be measured from the same well.

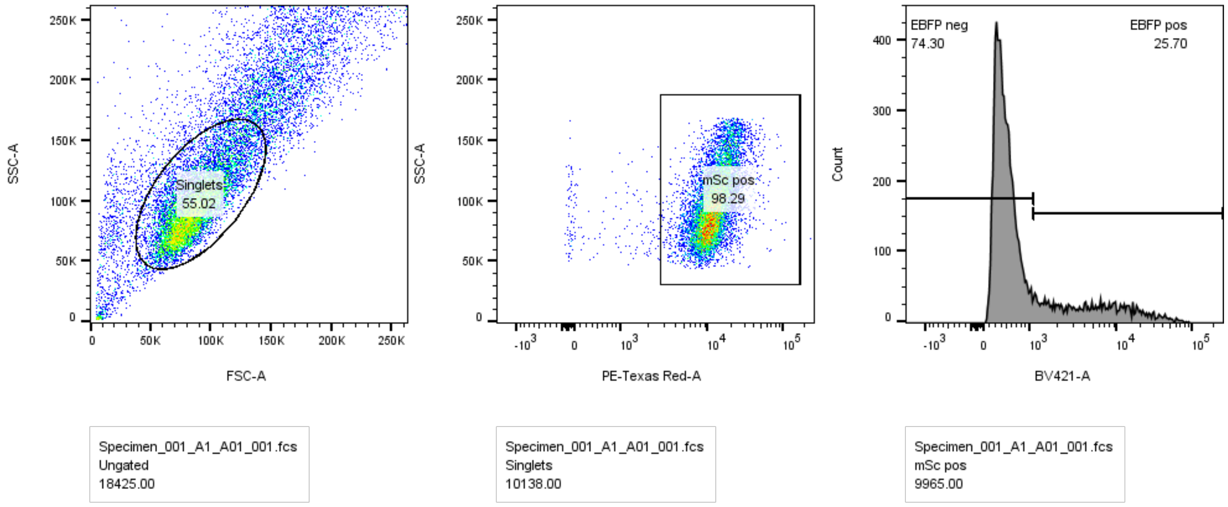

**Supplementary Fig. 15** | Example gating for the EBFP-gpASNase1 experiment.

#### Data analysis and statistics

Sample sizes, level of replication, summary statistic, and error bar types are stated in the figure captions. Independent experiment may refer to the same cell line but always refers to different experimental days (e.g. same experiment on same cell line but independently performed one week apart). Graphpad Prism 9 was used for statistical tests, using Welch ANOVA tests with Dunnett's T3 multiple comparison correction. P-values: 0.1234 (ns), 0.0332 (*), 0.0021 (**), 0.0002 (***), <0.0001 (****). Graphpad Prism 9 was used for curve fittings with the "Sigmoidal, 4PL, x is log(concentration)" equation (standard settings, no contraints or outlier elimination except where stated differently): Y=Bottom + (Top-Bottom)/(1+10^((LogIC50-X)*HillSlope))

#### iAspSnFR protein sequences

**"wild type" cell-surface screening construct (on pJTI vector)**

METDTLLLWVLLLWVPGSTGDRGS**AAGSTLDKIAKNGVIVVGHRESSVPFSYYDNQQKVVGYSQDYSNAIVE**
**AVKKKLNKPDLQVKLIPITAQNRIPLLQNGTFDFECGSTDNNVERQKQAAFSDTIFVVGTRLLTKKGGDIKDFA**
**NLKDKAVVVTSGTTSEVLLNKLNEEQKMNMRIISAKDHGDSFRTLESGRAVAFMMDDVLLAGERAKAKKPD**
**NWEIVGKPQSQEAYGCMLRKDDPQFKKLMDDTIAQVQTSGEAEKWFDKWFKNPILV**SHNVYITADKQKNGIK
ANFKIRHNVEDGSVQLADHYQQNTPIGDGPVLLPDNHYLSTQSVLSKDPNEKRDHMVLLEFVTAAGITLGMDEL
YKGGTGGSMSKGEELFTGVVPILVELDGDVNGHKFSVRGEGEGDATNGKLTLKFICTTGKLPVPWPTLVTTLT
YGVQCFSRYPDHMKQHDFFKSAMPEGYVQERTISFKDDGTYKTRAEVKFEGDTLVNRIELKGIDFKEDGNILG
HKLEYNFN**NPLNMNFELSDEMKALFKEPNDKALK**VDEQKLISEEDLNAVGQDTQEVIVVPHSLPFKVVVISAIL
ALVVLTIISLIILIMLWQKKPRGSKSRITSEGEYIPLDQIDINVMVSKGEAVIKEFMRFKVHMEGSMNGHEFEIEGE
GEGRPYEGTQTAKLKVTKGGPLPFSWDILSPQFMYGSRAFIKHPADIPDYYKQSFPEGFKWERVMNFEDGGA
VTVTQDTSLEDGTLIYKVKLRGTNFPPDGPVMQKKTMGWEASTERLYPEDGVLKGDIKMALRLKDGGRYLADF
KTTYKAKKPVQMPGAYNVDRKLDITSHNEDYTVVEQYERSEGRHSTGGMDELYKGSFCYENEV*

IgK signal peptide, **gltI binding protein,** cp-sfGFP, myc-epitope, PDGFR transmembrane domain, $K_{ir}$2.1 trafficking sequence, mScarlet-I, $K_{ir}$2.1 ER export motif, **underlined residues were diversified using deep mutational scanning**

**iAspSnFR for mammalian expression (on pcDNA3.1/Neo or pJTI vectors)**

MVSKGEAVIKEFMRFKVHMEGSMNGHEFEIEGEGEGRPYEGTQTAKLKVTKGGPLPFSWDILSPQFMYGSRA
FIKHPADIPDYYKQSFPEGFKWERVMNFEDGGAVTVTQDTSLEDGTLIYKVKLRGTNFPPDGPVMQKKTMGW
EASTERLYPEDGVLKGDIKMALRLKDGGRYLADFKTTYKAKKPVQMPGAYNVDRKLDITSHNEDYTVVEQYER
SEGRHSTGGMDELYKGS**AAGSTLDKIAKNGVIVVGHRESSVPFSYYDNQQKVVGFSQDYSNAIVEAVKKKLN**
**KPDLQVKLIPITAQNRIPLLQNGTFDFECGSTDNNVERQKQAAFSDTIFVVTTRLLTKKGGDIKDFANLKDKAV**

**VVTSGTTSEVLLNKLNEEQKMNMRIISAKDHGDSFRTLESGRAVAFMMDDVLLAGERARAKKPDNWEIVGK**
**PQSQEAWGCMLRKDDPQFKKLMDDTIAQVRTSGEAEKWFDKWFKNPILV**SHNVYITADKQKNGIKANFKIRH
NVEDGSVQLADHYQQNTPIGDGPVLLPDNHYLSTQSVLSKDPNEKRDHMVLLEFVTAAGITLGMDELYKGGTG
GSMSKGEELFTGVVPILVELDGDVNGHKFSVRGEGEGDATNGKLTLKFICTTGKLPVPWPTLVTTLTYGVQCF
SRYPDHMKQHDFFKSAMPEGYVQERTISFKDDGTYKTRAEVKFEGDTLVNRIELKGIDFKEDGNILGHKLEYNF
N**NPLNMNFELSDEMKALFKEPNDKALK***

**gltI binding protein,** cp-sfGFP, mScarlet-I, **underlined residues are mutations of iAspSnFR compared to SF-iGluSnFR S72A**

**EBFP-ASNase (on pSF-EF1a vector)**

MVSKGEELFTGVVPILVELDGDVNGHKFSVRGEGEGDATNGKLTLKFICTTGKLPVPWPTLVTTLSHGVQCFA
RYPDHMKQHDFFKSAMPEGYVQERTIFFKDDGTYKTRAEVKFEGDTLVNRIELKGVDFKEDGNILGHKLEYNF
NSHNIYIMAVKQKNGIKVNFKIRHNVEDGSVQLADHYQQNTPIGDGPVLLPDSHYLSTQSVLSKDPNEKRDHMV
LLEFRTAAGITLGMDELYKMARASGSERHLLLIYTGGTLGMQSKGGVLVPGPGLVTLLRTLPMFHDKEFAQAQ
GLPDHALALPPASHGPRVLYTVLECQPLLDSSDMTIDDWIRIAKIIERHYEQYQGFVVIHGTDTMASGASMLSFM
LENLHKPVILTGAQVPIRVLWNDARENLLGALLVAGQYIIPEVCLFMNSQLFRGNRVTKVDSQKFEAFCSPNLSP
LATVGADVTIAWDLVRKVKWKDPLVVHSNMEHDVALLRLYPGIPASLVRAFLQPPLKGVVLETFGSGNGPSKP
DLLQELRAAAQRGLIMVNCSQCLRGSVTPGYATSLAGANIVSGLDMTSEAALAKLSYVLGLPELSLERRQELLA
KDLRGEMTLPTADLHQSSPPGSTLGQGVARLFSLFGCQEEDSVQDAVMPSLALALAHAGELEALQALMELGS
DLRLKDSNGQTLLHVAARNGRDGVVTMLLHRGMDVNARDRDGLSPLLLAVQGRHRECIRLLRKAGACLSPQD
LKDAGTELCRLASRADMEGLQAWGQAGADLQQPGYDGRSALCVAEAAGNQEVLALLRNLALVGPEVPPAIGS
GSDYKDDDDK*

EBFP, gpASNase1, FLAG-tag

#### Supplemental references

1. Hellweg, L. *et al.* A general method for the development of multicolor biosensors with large dynamic ranges. 2022.11.29.518186 Preprint at https://doi.org/10.1101/2022.11.29.518186 (2022).
2. Craggs, T. D. Green fluorescent protein: structure, folding and chromophore maturation. *Chem. Soc. Rev.* **38**, 2865–2875 (2009).
3. Kabsch, W. XDS. *Acta Cryst D* **66**, 125–132 (2010).
4. McCoy, A. J. *et al.* Phaser crystallographic software. *J Appl Cryst* **40**, 658–674 (2007).
5. Emsley, P., Lohkamp, B., Scott, W. G. & Cowtan, K. Features and development of Coot. *Acta Cryst D* **66**, 486–501 (2010).
6. Murshudov, G. N. *et al.* REFMAC5 for the refinement of macromolecular crystal structures. *Acta Cryst D* **67**, 355–367 (2011).
7. Adams, P. D. *et al.* PHENIX: a comprehensive Python-based system for macromolecular structure solution. *Acta Cryst D* **66**, 213–221 (2010).

8. Chen, V. B. *et al.* MolProbity: all-atom structure validation for macromolecular crystallography. *Acta Cryst D* **66**, 12–21 (2010).
9. Karplus, P. A. & Diederichs, K. Linking Crystallographic Model and Data Quality. *Science* **336**, 1030–1033 (2012).
10. Girardot, C., Scholtalbers, J., Sauer, S., Su, S.-Y. & Furlong, E. E. M. Je, a versatile suite to handle multiplexed NGS libraries with unique molecular identifiers. *BMC Bioinformatics* **17**, 419 (2016).
11. Bushnell, B., Rood, J. & Singer, E. BBMerge – Accurate paired shotgun read merging via overlap. *PLOS ONE* **12**, e0185056 (2017).
12. Martin, M. Cutadapt removes adapter sequences from high-throughput sequencing reads. *EMBnet.journal* **17**, 10–12 (2011).
13. Hidalgo, F., Templeton, S., Gallegos, C. O. & Wang, J. Mutagenesis-Visualization: Analysis of Site-Saturation Mutagenesis Datasets in Python. 2021.10.08.463725 Preprint at https://doi.org/10.1101/2021.10.08.463725 (2021).
14. Gerst, R., Cseresnyés, Z. & Figge, M. T. JIPipe: visual batch processing for ImageJ. *Nat Methods* **20**, 168–169 (2023).
15. Stringer, C., Wang, T., Michaelos, M. & Pachitariu, M. Cellpose: a generalist algorithm for cellular segmentation. *Nat Methods* **18**, 100–106 (2021).
